# RNA silencing by CRISPR in plants does not require Cas13

**DOI:** 10.1101/2021.05.20.445036

**Authors:** VK Sharma, S Marla, WG Zheng, D Mishra, J Huang, W Zhang, GP Morris, DE Cook

## Abstract

RNA-targeting CRISPR-Cas can provide potential advantages over DNA editing, such as avoiding pleiotropic effects of genome editing, providing precise spatiotemporal regulation and expanded function including anti-viral immunity. Here, we report the use of CRISPR-Cas13 in plants to reduce both viral and endogenous RNA. Unexpectedly, we discovered that crRNA designed to guide Cas13 could, in the absence of the Cas13 protein, cause substantial reduction in RNA levels as well. We demonstrate Cas13-independent guide-induced gene silencing (GIGS) in three plant species, including stable transgenic *Arabidopsis*. We determined that GIGS utilizes endogenous RNAi machinery despite the fact that crRNA are unlike canonical triggers of RNAi such as miRNA, hairpins or long double-stranded RNA. These results suggest that GIGS offers a novel and flexible approach to RNA reduction with potential benefits over existing technologies for crop improvement. Our results demonstrate that GIGS is active across a range of plant species, evidence similar to recent findings in an insect system, which suggests that GIGS is potentially active across many eukaryotes.

## Introduction

Genome editing technologies such as CRISPR-Cas9 (clustered regularly interspaced short palindromic repeats and CRISPR associated protein), CRISPR-Cas12, and newly identified systems, enable unprecedented opportunities for genome engineering ^1–4^. However, DNA editing technologies involving double-strand break repair can result in the creation of unintended DNA mutations^5,6^, potentially hindering applications. The derivative Cas9 protein, termed PRIME-editor, enables more precise editing and overcomes the unintended consequences resulting from the creation of double-strand breaks ^7^. Despite these technical advances in genome engineering, there remains a potentially fundamental limitation to DNA editing, where the alteration of a gene results in unintended and unpredictable phenotypes. This will occur for genes with pleiotropic effects ^8^. Additionally, many target traits for improvement are polygenic in nature, and multi-gene genome editing will compound the problem of generating unwanted phenotypes^9^. One approach to overcome these limitations is spatiotemporally genome editing, such as demonstrated with the CRISPR tissue-specific knockout system (CRISPR-TSKO), in which DNA is edited in specific cell types^10^. This approach will likely serve a role in future application of genome engineering, but the generation of mosaic genotypes caused by differences in the rate and penetrance of cell-specific editing, especially in polyploid crops, may limit the utility of this approach.

An alternative approach is the manipulation of RNA as it plays a central role in cellular dynamics, mediating genotype-phenotype relationship in eukaryotes. Manipulating RNA has potential advantages over DNA editing, such as circumventing negative pleiotropy, where an RNA product can be specifically spatiotemporally regulated. To manipulate complex traits, the targeting of multi-copy genes or multi-gene pathways through RNA manipulation offers more flexibility and precision than DNA editing approaches. Further, RNA manipulation can also be used to target RNA viruses for engineered immunity ^11^. Current RNA degradation technologies involving RNA interference (RNAi) suffer from off-target silencing ^12^, potentially introducing the same pleiotropic and unintended phenotypes as DNA editing.

To overcome these limitations, we sought to develop the class II type VI CRISPR-Cas13 system for use in plants, where the Cas13 nuclease specifically binds target single-stranded (ss)RNA in a CRISPR RNA (crRNA) guided manner ^13–15^. Recent reports have established the use of Cas13 as an introduced anti-viral immune system in plants ^16–18^. Here we report the discovery that crRNA guides alone, in the absence of Cas13, cause the reduction of both viral and endogenous plant mRNA in a sequence dependent manner. Mechanistically, our results suggest this guide-induced gene silencing (GIGS) functions through endogenous components of the RNAi pathway and are dependent on Argonaute protein(s). The use of compact, multi-guide crRNA to elicit selective RNA reduction provides a new avenue, along with Cas13-dependent approaches, to precisely manipulate plant traits.

## Results

### crRNA guides alone, in the absence of Cas13, can elicit target RNA reduction

To test the Cas13 system in plants, we synthesized the coding sequence for two Cas13a proteins, termed LbaCas13a (from *Lachnospiraceae bacterium*) and LbuCas13a (*Leptotrichia buccalis*) for expression in plants. We tested their function *in planta* by targeting the plant infecting Turnip mosaic virus (TuMV) expressing GFP by co-expressing Cas13, crRNA targeting TuMV, and TuMV expressing GFP in *Nicotiana benthamiana* leaves using *Agrobacterium-*mediated transient expression^19,20^. The Cas13 proteins were expressed with a single-guide crRNA containing antisense sequence to one region of the TuMV genome (single-guide), a multi-guide crRNA containing sequence against three regions of the genome (multi-guide), or an empty-guide, which contained the direct repeat (DR) crRNA sequence alone (Fig. 1a). Expression of either Cas13a protein with the single- or multi-guide crRNA reduced viral accumulation by 72 hours post inoculation (hpi) (Supplementary Fig. 1a). Virus accumulation was reduced by approximately 90% at 120 hpi, and TuMV interference by Cas13a was dependent on the expression of a crRNA with complementary sequence (Supplementary Fig. 1b-d).

**Figure 1.**
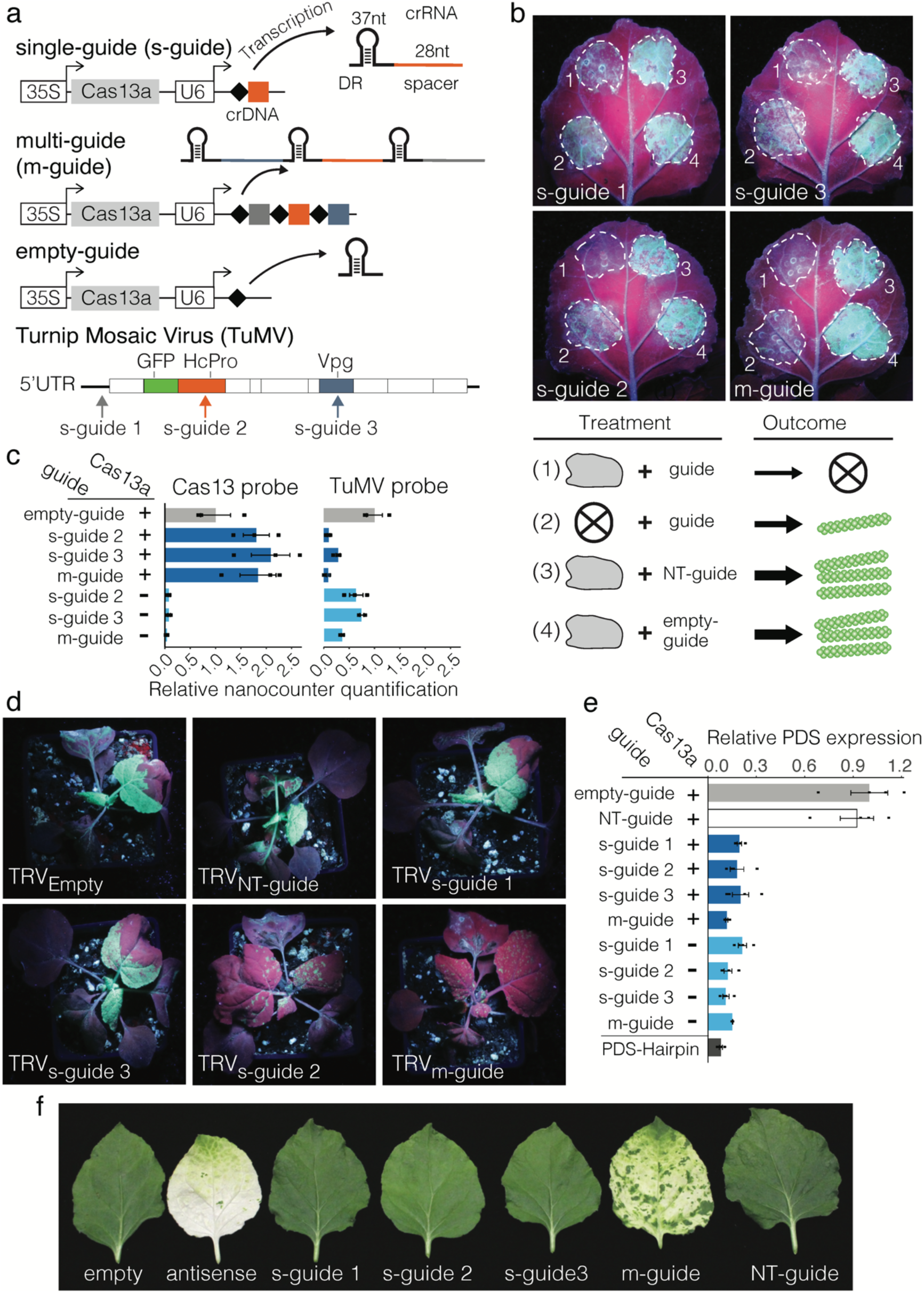
Cas13 and GIGS reduce viral and endogenous target RNA in *N. benthamiana*. **a,** Schematic overview of the Cas13 transgene system. Guide crRNA responsible for RNA target specificity contain a single 28 nucleotide (nt) spacer antisense to the target RNA (single-guide, s-guide), multiple 28 nt spacers (multi-guide, m-guide), or lack the spacer (empty-guide). A diagram showing the genome of turnip mosaic virus (TuMV) expressing GFP and indicating the location the three targeting sites for the guide crRNA. **b**, The accumulation of GFP was assessed at 120 hours post inoculation based on GFP fluorescence. Areas of agroinfiltration are shown in dashed white circles. Individual treatments are labeled with numbers and shown schematically below the photographs. **c**, Nanostring RNA quantification for Cas13 and TuMV levels corresponding to labeled treatments for *N. benthamiana* spot infiltration. Samples expressed Cas13 (+) or not (-). **d**, Representative images of *N. benthamiana* plants under UV light at 7 days post inoculation. The systemic movement of TuMV is evident based on the accumulation of GFP fluorescence for empty-guide expressing TRV (TRV_empty_). Single-guide 2 and multi-guide, TRV_s-guide 2_, and TRV_m-guide_ respectively, stopped systemic TuMV infection. **e**, Quantitative PCR for the endogenous transcript *PDS* following *N. benthamiana* leaf spot infiltration. **f**, Representative single leaf images of *N. benthamiana* following TRV-mediated systemic delivery of guide crRNA targeting the *PDS* transcript. Empty and non-target guides (NT-guide) did not cause photobleaching (white sectors), while the antisense and multi-guide (m-guide) did induce visible photobleaching.

In CRISPR-Cas experiments, the negative control characterizing cells expressing the sgRNA or crRNA alone, without Cas, are generally omitted due to the assumption of Cas-dependence. Interestingly, we observed that expression of a single-guide or multi-guide crRNA alone, in the absence of the Cas13a protein, inhibited viral accumulation as evidenced by reduced viral genome and derived protein accumulation (Fig. 1b and Supplementary Fig. 2a). Viral RNA was also directly quantified using two independent NanoString nCounter probes, which allowed direct RNA quantification without the creation of complementary (c)DNA. Probes against two different regions of the TuMV genome confirmed that the single-guide and multi-guide caused virus interference when expressed with Cas13a, but also when expressed alone, in the absence of Cas13a (Fig. 1c and Supplementary Fig. 2b). The NanoString quantification indicated that LbuCas13a plus guides provided greater viral interference compared to the single- or multi-guide alone. Among the samples expressing guide crRNA alone, the multi-guide consistently caused the greatest TuMV reduction compared to the single-guides (Fig. 1b,c and Supplementary Fig. 2a,b)

To determine whether GIGS can function systemically, GIGS-mediated TuMV interference was tested using the tobacco rattle virus (TRV) expression system^21^. Plants were co-inoculated with TuMV expressing GFP and TRV, which systemically produced single- and multi-guide crRNA in the absence of Cas13 (Supplementary Fig. 3a). At 7 days post inoculation (dpi), GFP-fluorescence from TuMV was observed in the upper systemic leaves of plants co-inoculated with either TRV expressing an empty-guide or a non-targeting (NT)-guide, which showed that systemic TRV delivery alone did not interfere with TuMV replication, movement, or translation (Fig. 1d). Samples expressing the two single-guides, s-guide 1 and s-guide 3, also accumulated visible GFP fluorescence in upper, non-inoculated leaves, indicating the spread of TuMV. Interestingly however, TRV expressing either single-guide 2 or the multi-guide caused a significant reduction in GFP-fluorescence in the upper systemic leaves (Fig. 1d, and Supplementary Fig. 3b). Quantitative assessment of TuMV accumulation in systemic leaves by qPCR showed an approximately 90% reduction in TuMV accumulation in samples expressing single-guide 2 and the multi-guide (i.e. GIGS) (Supplementary Fig. 3c). Moreover, qPCR revealed an approximate 30% to 40% reduction in TuMV levels when TRV expressed single-guide 1 or - guide 3, which was not obvious from visual inspection of GFP fluorescence. This may reflect complicated translation mechanisms viruses employ, such as internal ribosome entry^22^, in which the viral molecule was targeted by GIGS and partially interfered with, while intact GFP open reading frame sequence was still translated. These results indicate that GIGS can cause systemic TuMV interference, but that crRNA target sequences vary in effectiveness. Variation for crRNA effectiveness has been reported for Cas13-dependent RNA targeting, likely caused by secondary structure and accessibility of the target RNA^23^.

Viruses manipulate host physiology and have unique features unlike host derived RNAs^24,25^, making it possible that the GIGS phenomena is limited to viral RNA. To test this hypothesis, we targeted endogenous phytoene desaturase (*PDS*) mRNA with single-guide and multi-guide crRNA with and without LbuCas13a (Supplementary Fig. 4). *Agrobacterium*-mediated expression of single- and multi-guide crRNA with and without LbuCas13 caused a significant reduction in *PDS* transcript levels compared to expressing LbuCas13a alone or with a NT-guide (Fig. 1e). The resulting mRNA reduction (75-85%) was consistent across the tested samples, comparable to a PDS-hairpin construct known to induce RNAi (Fig. 1e). The reduction in PDS mRNA was confirmed by northern blot, which showed a clear reduction for PDS signal for both LbuCas13a-dependent and GIGS compared to expressing LbuCas13a alone, with a NT-guide, or from an untreated leaf (Supplementary Fig. 5a). Direct RNA quantification by NanoString further confirmed a significant reduction for the *PDS* transcript for samples expressing the *PDS* targeting guides with or without the expression of Cas13a (Supplementary Fig. 5b). These results establish that GIGS acts on both viral RNA and endogenous transcripts.

To test if GIGS acts systemically on endogenous genes, TRV expressing guides targeting endogenous *PDS* mRNA were infiltrated into *N. benthamiana* (Supplementary Fig. 6). Under the hypothesis that GIGS can act systemically on endogenous genes, the prediction is that TRV-delivered guides result in photobleaching in TRV-infected tissues. Three single-guide crRNA, targeting different regions of *PDS*, did not exhibit significant photobleaching (Fig. 1f). However, two multi-guides targeting different *PDS* regions displayed substantial photobleaching in systemic leaf tissue (Fig. 1f and Supplementary Fig. 7a). Interestingly, the visible photobleaching pattern induced by the anti-sense fragment (i.e. RNAi) and that induced by GIGS were not the same (Fig. 1f and Supplementary Fig. 7a). While the antisense RNAi photobleaching was strong in the upper, youngest leaves, GIGS induced photobleaching was not visible in the upper most leaves, and the photobleaching occurred in more distinct segments causing a patchy appearance. Quantifying the photobleaching to confirm the phenomena, SPAD meter readings showed a significant reduction in chlorophyll content for samples expressing the multi-guide crRNAs and containing the antisense PDS fragment (Supplementary Fig. 7b). Plants that expressed single-guide 2 were yellow and also showed a reduced SPAD reading (Supplementary Fig. 7a,b). Quantifying *PDS* transcripts with qPCR showed that the *PDS* transcript level was reduced (30-45%) for the three single-guides, and to a greater extent by the multi-guides (65-70%) and the antisense construct (85%) (Supplementary Fig. 7c). It is not clear why single-guide 1 and 3 caused a reduction in *PDS* mRNA levels, but did not result in visible photobleaching or SPAD meter reductions, but we note that the reduced *PDS* mRNA levels are consistent with that seen using *Agrobacterium*-mediated spot infiltration (e.g. Fig. 1e and Supplementary Fig. 5). Collectively, we found that GIGS induced by multi-guides caused a greater reduction in target transcript levels compared to that induced by single-guides for both virus and endogenous RNA targeting.

### GIGS functions in multiple plant species and is heritable in *Arabidopsis*

An important question is whether GIGS is limited to *N. benthamiana* or is more broadly active in plants. To test this, multi-guide crRNA were developed to target *PDS* in tomato (*Solanum lycopersicum*), which were delivered using TRV, along with a NT-guide and an antisense PDS control. We observed visible photobleaching in upper leaves of *S. lycopersicum* plants following systemic movement of TRV expressing a multi-guide targeting *S. lycopersicum PDS*, although the photobleaching was not as widespread as that produced by the antisense PDS construct (Fig. 2a). Quantifying chlorophyll levels and the *PDS* transcript indicated that photobleached tissue from GIGS and antisense expressing TRV both had substantially lower levels compared to the control (Fig. 2b,c). These results show that GIGS is active outside of *N. benthamiana*, possibly extending to other plants in the *Solanaceae* family.

**Figure 2.**
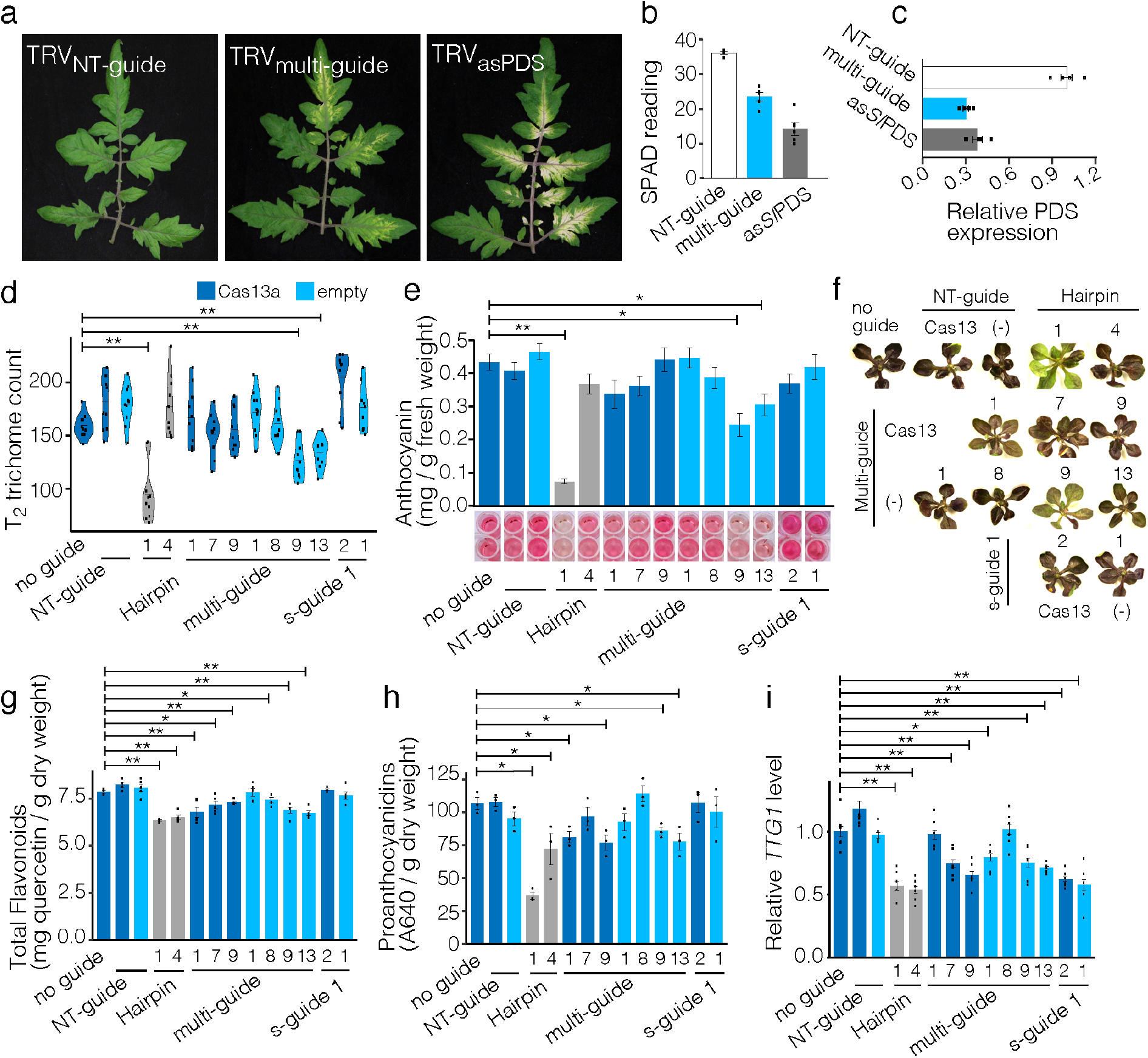
Cas13 and GIGS function across plant species and are heritable. **a**, Representative images of tomato leaves following TRV systemic movement and photobleaching induced by GIGS (TRV_m-guide_) and an antisense transcript (TRV_asPDS_). TRV expressing a non-targeting guide crRNA (TRV_NT-guide_) does not induce photobleaching. **b**, Measurements of chlorophyll content from SPAD meter readings for three independent plants. SPAD meter readings were taken from leaf sections showing photobleaching, and individual reading are shown as black points with the mean and standard deviation shown as a bar plot. **c**, qPCR measurement of the *PDS* transcript standardized to the *EF1*α transcript and relative to the NT-guide sample. Three independent samples were analyzed and individual data are shown as black points with the mean and standard deviation shown as bar plots. (**d-i)**, Data for independent transgenic *Arabidopsis* lines. Data for plants expressing LbuCas13a are shown in dark blue and plants not expressing the protein are shown in light blue. Control lines expressing a hairpin construct against the *TTG1* transcript are shown in grey. **d**, Trichome counts from the seventh leaf of T_2_ *Arabidopsis* lines. Ten plants were counted per independent line, listed below graph, with the individual counts shown as black points and the distribution represented as a violin plot. **e**, Leaf anthocyanin quantification from T_3_ seedlings following sucrose treatment. Representative wells follow extraction shown below each bar plot. **f**, Representative plantlets following sucrose treatment showing anthocyanin pigmentation (i.e. purple color). **g**, Total flavonoids extracted from seeds collected from T_2_ plants. Five independent seeds lots were analyzed per line, shown as black points. **h**, Seed proanthocyandin quantification from the same plants analyzed in (g). **i**, Quantification of the *TTG1* transcript from three T_2_ and three T_3_ plants per line, individual data shown as black points. Statistical comparisons were made between the transformation control (no guide) and each treatment using a one-sided Mann-Whitney U-test with Benjamini-Hochberg (BH) multiple testing correction. Samples with p-values less than 0.05 (*), and 0.01 (**) are indicated.

Another important question is whether GIGS requires bacterial or viral machinery (i.e. proteins) introduced during transient expression or if GIGS functions in stable transgenics through plant endogenous machinery. To test these hypotheses, and further test the generality of GIGS in plants, we transformed *Arabidopsis thaliana* (Col-0) with single-guide and multi-guide crRNA targeting the pleiotropic regulatory gene *TRANSPARENT TESTA GLABRA1* (*TTG1*), both with and without LbuCas13a. The *TTG1* gene encodes a WD40 repeat protein, which interacts with MYB and bHLH transcription factors required for normal trichome and root hair development, along with seed proanthocyanidin and vegetative anthocyanin production^26–28^. The average trichome counts for multiple independent T_1_ plants that expressed LbuCas13a with either single-guide or multi-guide crRNA had significantly fewer trichomes compared to wild-type, and importantly, plants expressing single-guides and the multi-guide crRNA, without Cas13, also had significantly fewer trichomes on average (Supplementary Fig. 8a). The *TTG1* transcript was quantified in T_1_ plants and was highly variable across the transformed lines (Supplementary Fig. 8b). Individual plants were selected, self-fertilized and seeds from T_1_ plants showed reduced total flavonoids in both Cas13 and GIGS lines, consistent with reduced *TTG1* (Supplementary Fig. 8c).

We assessed whether GIGS would function in progeny inheriting guides by characterizing individual lines in the T_2_ and T_3_ generations for alteration of *TTG1*-dependent phenotypes. Trichome counts of the seventh leaf (from ten plants per line) indicated that two GIGS lines (i.e. expressing only a multi-guide crRNA targeting *TTG1*), and one of the hairpin expressing lines had significantly fewer trichomes compared to the transformation control expressing Cas13a alone (Fig. 2d). Individual transformed lines were subjected to sucrose and light stress to induce leaf anthocyanin production, and we again observed that two lines expressing multi-guide crRNA targeting *TTG1* (i.e. GIGS) displayed significantly reduced leaf anthocyanin levels, along with a hairpin expressing line (Fig. 2e,f). Quantification of total seed flavonoids showed a significant but modest reduction compared to the control line, for both Cas13 expressing and GIGS lines along with both hairpin expressing lines (Fig. 2g). Total flavonoid quantification also measures products upstream of *TTG1* regulation, which can confound the impact of *TTG1* reduction. To more accurately assess the impact of *TTG1* reduction, we measured seed proanthocyanidins, which are controlled downstream of *TTG1*. This analysis identified a more substantial impact for *TTG1* reduction, where the level of proanthocyanidins were significantly reduced (Fig. 2h), and were consistent with the results from the total flavonoid quantification (Fig. 2g).

These results indicate heritable phenotypes for multiple traits mediated by both Cas13 and GIGS in stable transgenic *Arabidopsis* when targeting the pleiotropic regulator *TTG1*. We do note there was substantial phenotypic variation among lines with the same construct, despite significant reduction in *TTG1* levels (Fig. 2i). This is in part explained by variation in transgene expression and translation (Supplementary Fig. 9). In addition, more complicated mechanisms such as asynchronous *TTG1* expression and Cas13 or GIGS expression at the individual cell level, or the effect of incomplete *TTG1* silencing on trait manifestation (i.e. kinetics of silencing to produce a phenotype)^29,30^. Optimizing Cas13 and GIGS approaches will be an important step to deliver robust biotechnology platforms for plant research and crop improvement, particularly for tissue- or temporal-specific expression that is difficult to manipulate precisely with CRISPR-Cas9.

### Multi-guide crRNA induce secondary small RNA production

We sought to understand the mechanism giving rise to GIGS (i.e. guide crRNA reducing viral and endogenous RNA levels). Given that crRNA are composed of short antisense sequences, it is possible that GIGS functions through components of the endogenous RNA interference (RNAi) pathway. However, the structure of crRNA used here are not similar to hairpin RNA, small interfering RNA (siRNA), or micro RNA (miRNA), therefore it is not obvious how crRNA might enter or induce RNAi^31,32^. Alternatively, it is possible that GIGS elicits other endogenous endo- or exonucleolytic RNA degradation pathways^33^. Since small RNA (sRNA) usually in the range of 21-to 24-nucleotides (nt) are a hallmark for RNAi, we reasoned that if GIGS functions through RNAi, abundant sRNA should be observed^34^. To assess this, we conducted small (s)RNA-seq from *N. benthamiana* samples expressing single and multi-guide crRNA against the endogenous *PDS* transcript. Uniquely mapped sRNA for the single-guide samples showed a single sharp peak at the *PDS* transcript, which corresponds to the location of the crRNA guide sequence, regardless of Cas13 expression (Fig. 3a). Likewise, the samples expressing the multi-guide crRNA had three distinct peaks of mapped sRNA, each corresponding to the location of the targeting guide sequence. However, in these samples we also identified many sRNA mapping to the *PDS* transcript that were independent from the multi-guide target sequence (Fig. 3b). Interestingly, these sRNA were identified only between the 5’ and 3’ boundaries of crRNA targeting sites and do not appear to extend past this region (Fig. 3b). This was similar to the sRNA mapping from the samples expressing the PDS hairpin, which produced ample sRNA between the two ends of the hairpin fragment (Fig. 3c). While the most abundant peaks for the multi-guide crRNA samples corresponded to the guide targets themselves, the identification of thousands of sRNA reads between these target regions suggest the production of secondary sRNA. We do note the presence of background sRNA in the samples where Cas13 was expressed with a NT-guide, which may indicate background read mapping or potentially RNA contamination during library preparation, but the signal was low (Fig. 3d). Supporting the idea that GIGS results in the production of secondary sRNA through RNAi, we identified more 21 nt sRNA (i.e. siRNA) mapped to the *PDS* transcript during GIGS (i.e. without the Cas13 protein) than when Cas13 was expressed with the guide (Fig. 3e).

**Figure 3.**
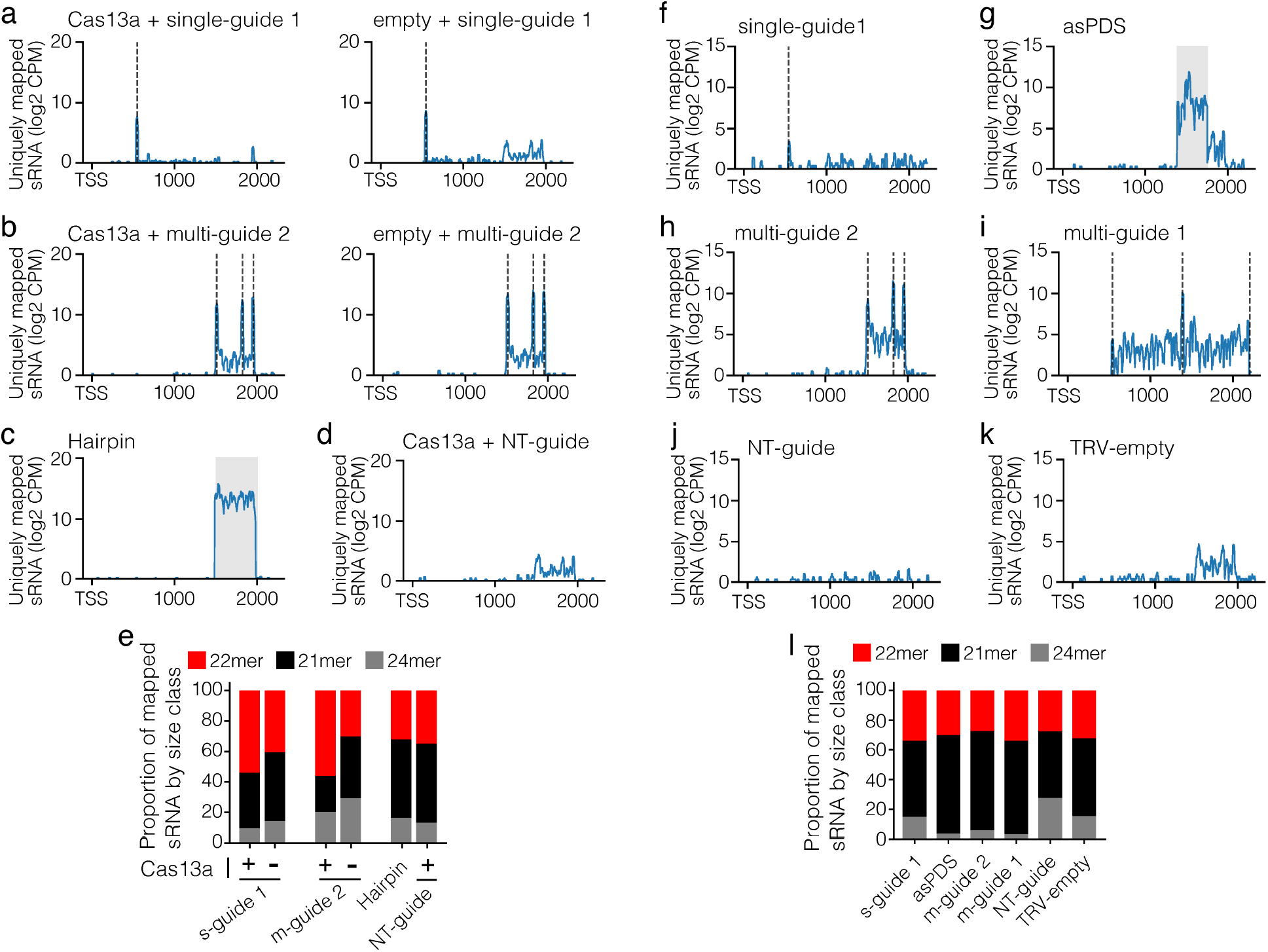
Multi-guide induced GIGS results in sRNA generation. (**a-d**), Uniquely mapped small RNA (sRNA) read counts to the *PDS* transcript collected five days post agro-mediated spot infiltration. Read counts are log_2_ of counts per million +1 (log_2_ CPM) and shown relative to the transcription start site (TSS) till the end of the predicted mRNA (2216 bp). Individual treatments are labeled above each graph, and one of the two replicate samples per treatment is plotted. The position of the expressed single- and multi-guide crRNA are shown as vertical dashed line(s). The region spanning the hairpin construct is shown as a grey window. (**e**), Proportion of 21-, 22-, and 24-nt sRNA mapped to the *PDS* transcript averaged between the two replicates. (**f-l**), similar layout as described in (a-e) but here RNA was collected from systemic leaves two-weeks following TRV expression. The treatments are listed above each graph.

To further determine sRNA production during GIGS, a second sRNA-seq experiment was conducted by expressing either a single-guide or one of two multi-guide crRNA in the absence of Cas13 using the TRV vector in *N. benthamiana*. The uniquely mapped sRNA from the single-guide had a clear but small peak corresponding to the guide target sequence, along with other background mapped sRNA (Fig. 3f). In contrast, mapped sRNA from the sample expressing a PDS antisense fragment produced many sRNA, which mapped between the ends of the antisense fragment (Fig. 3g). Both multi-guide crRNAs showed three sharp peaks of mapped sRNA, with each peak corresponding to a guide targeting region (Fig. 3h,i). Importantly, these samples clearly have many mapped sRNA that are outside of the multi-guide targeted region, which are not present in the controls, and were not expressed as part of the multi-guide crRNA sequence (Fig. 3h-k). We interpret these sRNA to represent secondary sRNA generated in response to multi-guide GIGS. Consistent with these secondary sRNA being generated via components of the RNAi pathway, the length of sRNA mapped to the *PDS* transcript are predominantly 21 nt for the two multi-guide and antisense fragment samples (Fig. 3l). These results suggest that siRNA and RNAi are likely involved in mediating GIGS.

### GIGS RNA reduction functions through Argonaute

Under the hypothesis that GIGS requires endogenous RNAi machinery, target mRNA reduction would be dependent on Argonaute (AGO) RNA-binding protein(s)^35^. AGO proteins are required to form the RNA Induced Silencing Complex (RISC), which carries out the biochemical slicing or translational inhibition of target mRNA^36,37^. To achieve AGO mediated endonuclease activity, perfect complementary base pairing is required at positions 10 and 11 of the AGO-bound siRNA with the target mRNA (i.e. central duplex region)^38–40^. Therefore, if GIGS is dependent on AGO, multi-guide crRNA designed to have mismatches at base-pairs 10 and 11 should be blocked for GIGS (i.e. no target mRNA reduction). To test this, multi-guide crRNA that contained specific two base pair mismatches to the PDS mRNA were delivered to *N. benthamiana* using TRV (Fig. 4a). The results showed that multi-guide crRNA against *PDS* with mismatches at the critical region for AGO endonuclease activity (i.e. base pairs 10,11) did not cause photobleaching, while negative control mismatches (i.e. positions 5,6 or 21,22) still elicited photobleaching (Fig. 4a, Supplemental Fig. 10 for whole plant images). The chlorophyll content as measured by SPAD meter was not significantly different between the NT-guide control and the multi-guide with mismatches at positions 10,11 (mg 1[mm10,11]) (Fig. 4c). The perfect complementary multi-guide, along with the guide containing mismatches at positions 5,6 and 21,22 had significantly reduced SPAD meter readings, along with the antisense PDS construct (Fig. 4c). Quantification of *PDS* transcripts by qPCR confirmed no reduction for samples expressing the multi-guide with position 10,11 mismatches, while all other treatments significantly reduced the level of the *PDS* transcript (Fig. 4d). We note that the mismatches at 5,6 and 21,22 did affect silencing, as the perfectly complementary multi-guide crRNA gave the strongest photobleaching. These mismatches may interfere with other RISC functions, such as target recognition and target mRNA turnover^38,40^. However, it is clear that mismatches at 10,11 abolish GIGS, while the other mismatches diminish it, suggesting that GIGS functions through one or more endogenous AGO proteins. Additionally, these results suggest that GIGS is mediated by RNA endonuclease reduction and not translational inhibition of target mRNA^41^.

**Figure 4.**
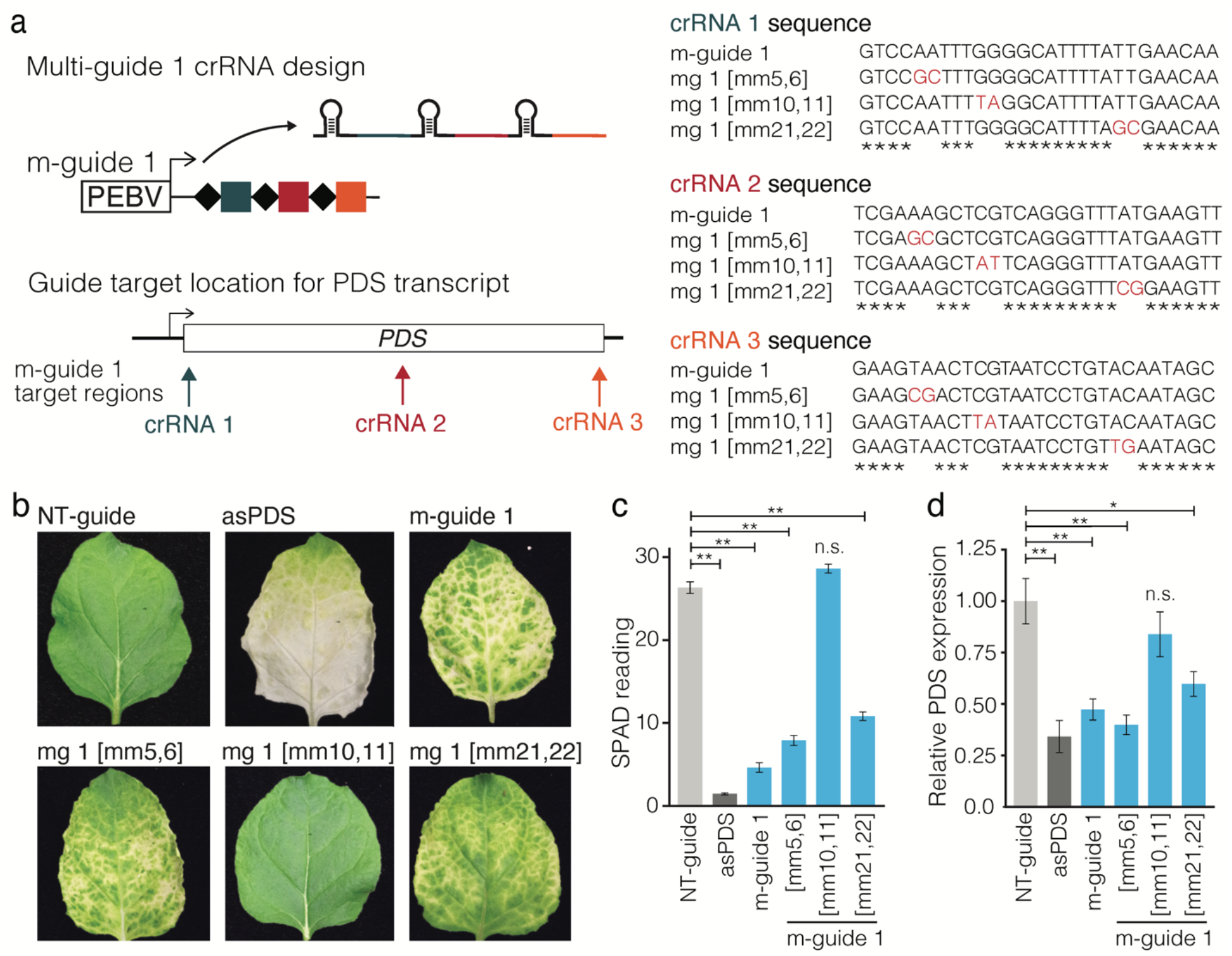
Guide mismatches at position 10 and 11 abolish GIGS, indicating AGO dependence. (**a**), Illustration of multi-guide expression from TRV targeting the *PDS* transcript. For each of the 28 nt guides (crRNA1, crRNA2, crRNA 3) a variant m-guide 1 was designed. For mg 1[mm5,6], each crRNA contained two base pair mismatches at positions 5,6, for mg 1[mm10,11] mismatches at positions 10,11, and mg 1[mm21,22] contained mismatches at positions 21,22. (**b**), Representative images of leaves following TRV systemic delivery of m-guide 1 targeting PDS, in addition to the three variants of m-guide 1. TRV expressing a non-targeting guide (NT-guide) and TRV with a region of antisense sequence to PDS (asPDS) served as controls. (**c**), SPAD meter readings from photobleached (loss of green color) leaf samples. Data collected from a total of six independent leaves from two experiments. (**d**), Quantification of the *PDS* transcript using qPCR for the same samples as measured in (c). Data standardized to an endogenous transcript and normalized to TRV expressing NT-guide. Statistical comparisons were made between the NT-guide and each treatment using a one-sided Mann-Whitney U-test with Benjamini-Hochberg (BH) multiple testing correction. Samples with p-values less than 0.05 (*), and 0.01 (**) are indicated. n.s., non-significant difference (p > 0.05).

### GIGS also occurs with Cas9 designed crRNA

The Cas13 guide crRNA are composed of the Cas13 specific direct repeat (DR) domain and the antisense target sequence ^42^, and they do not contain double-stranded RNA corresponding to the target sequence as would be found in a hairpin, short-hairpin or miRNA transgene. It was therefore not clear if a sequence or structure of Cas13 designed crRNA were required to elicit GIGS. It was recently reported that crRNA guides from the Cas13b system cause target mRNA reduction in the absence of Cas13b, termed Cas13b-independent silencing in mosquito ^43^. That report does not provide functional data that elucidate the mechanism, but the authors postulate that Cas13b-independent silencing is related to RNAi. Importantly, the Cas13b DR sequence is different than the Cas13a DR sequence used here. Additionally, the structure of the crRNA are different, where the Cas13b DR is located at the 3’ end of the crRNA following the target guide sequence, while the Cas13a crRNA used here have a 5’ DR prior to the target sequence ^42^. These results suggest that GIGS is not dependent on a specific Cas13 DR sequence or structure. To directly investigate this hypothesis, we tested if GIGS was active for other guide crRNA, such as for the CRISPR-Cas9 system. Using the Cas13 single-guide (s-guide 2) that caused a slight yellowing in the leaf and *PDS* mRNA reduction (Fig. 1d,e), we designed a corresponding 28 nt Cas9 sgRNA (Fig. 5a, sgRNA 1). When the Cas9 designed sgRNA was delivered by TRV, we observed subtle yellowing in the leaves compared to TRV expressing a NT-guide, similar to that produced by the Cas13 crRNA design (Fig. 5b). Importantly, a control Cas9 designed sgRNA containing 50% mismatches to the PDS sequence showed no yellowing, indicating that the subtle phenotype was specific (Fig. 5b and Supplementary Fig. 11 for whole plant images). This visible phenotype was corroborated by SPAD meter readings that indicated an approximately 28% reduction in chlorophyll content compared to the control expressing a NT-guide, which was similar to the reduction observed for the Cas13 designed s-guide (Fig. 5c). Molecular quantification indicated significant but variable *PDS* transcript reduction compared to the NT-guide and the 50% mismatch sgRNA controls (Fig. 5d).

**Figure 5.**
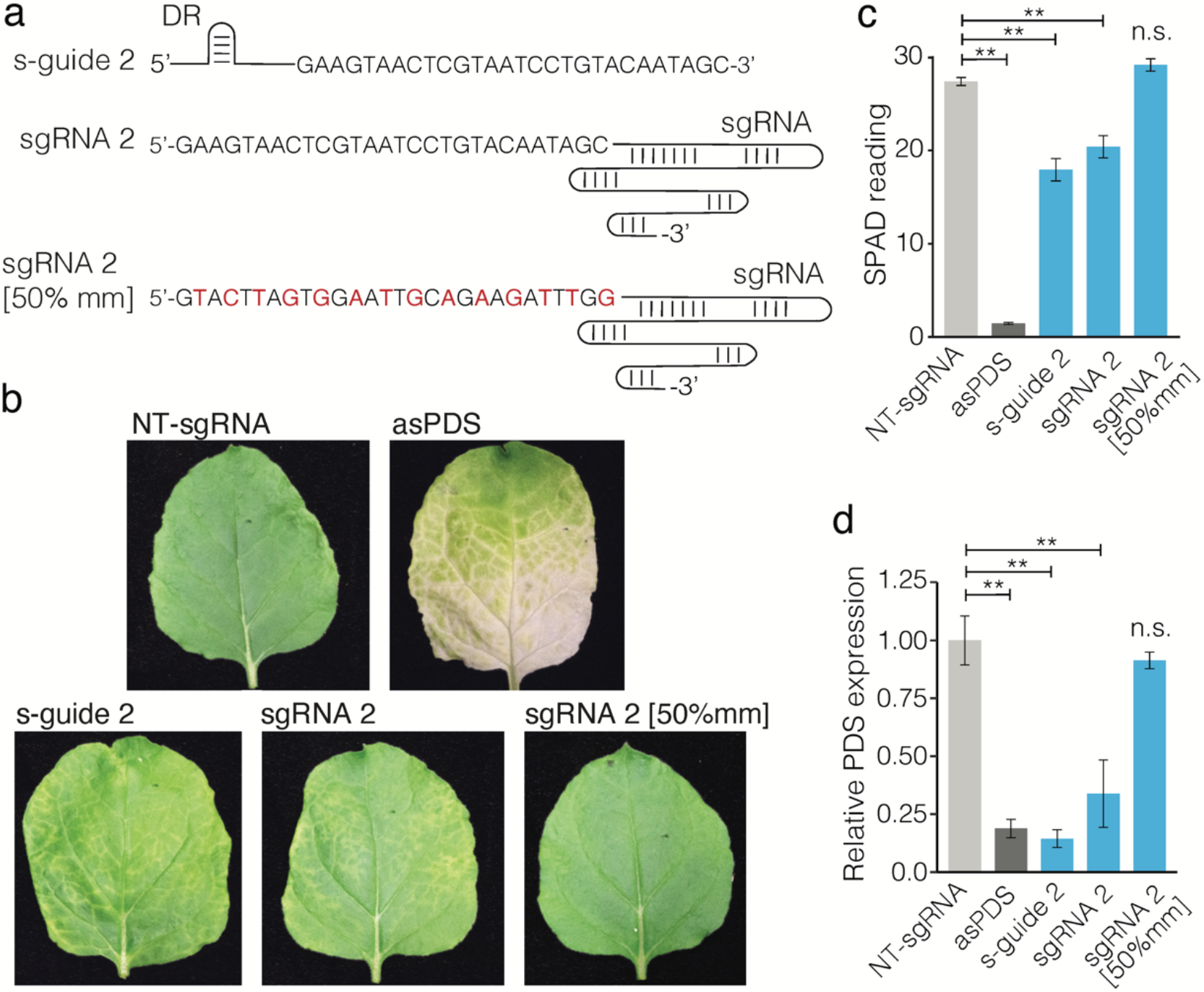
GIGS is also evident for sgRNA guides designed for Cas9. (**a**), Schematic of guide designs targeting *PDS* transcript for Cas13 s-guide 2, and Cas9 sgRNA 2. Each guide contains 28 nt antisense to the *PDS* transcript (sequence shown). The Cas9 sgRNA control contained 50% mismatch sequence to the *PDS* transcript (sgRNA 2 [50% mm]). Mismatch nucleotides are colored red, while shared nucleotides between the three guides are black. The Cas13 crRNA contains the 37 bp direct repeat (DR) sequence at the 5’ end. The Cas9 sgRNA contains the 78 bp trans-activating crRNA (tracrRNA, depicted as a line) at the 3’ end. (**b**), Representative images of leaves following TRV systemic delivery of single-guide 2 (s-guide 2) targeting *PDS*, and a Cas9 designed sgRNA designed to contain the same 28 bp targeting PDS as in s-guide 2. An sgRNA 2 control contained the sequence in sgRNA 2, but with 50% mismatches to the *PDS* transcript (sgRNA 2[50%mm]). Control sgRNA containing non-targeting guide sequence (NT-sgRNA). Photobleaching is seen in the asPDS sample, while interveinal yellowing is visible in the samples expressing s-guide 2 and sgRNA 2. (**c**), SPAD meter readings from photobleached leaf samples as described for (b). Data collected from a total of six independent leaves from two experiments. (**d**) Quantification of the *PDS* transcript using qPCR for the same samples as measured in (c). Data standardized to an endogenous transcript and normalized to TRV expressing NT-sgRNA. Statistical comparisons were made between the NT-sgRNA and each treatment using a one-sided Mann-Whitney U-test with Benjamini-Hochberg (BH) multiple testing correction. Samples with p-values less than 0.05 (*), and 0.01 (**) are indicated. n.s., non-significant difference (p > 0.05).

## Discussion

The rapid pace of biotechnological innovation for trait manipulation is advancing science and has incredible potential to benefit society. CRISPR-based approaches for RNA manipulation offer new approaches for trait manipulation, but they are currently less well understood compared to DNA-targeting CRISPR. Through the course of our work to develop Cas13 for use in plants, we unexpectedly discovered that the guide crRNA designed for the Cas13a system can reduce viral and endogenous RNA in the absence of the Cas13 protein (i.e. GIGS). There is a question of why this was not previously reported in plants. One explanation is that previous reports of Cas13 function in plants and other systems have not included a guide-alone control (e.g. stable transgenic line expressing guide crRNA alone) such as the experiments described for stable transgenic rice 17, rice protoplasts^15^, and experiments in animal systems^15,44,45^. One experiment did test for guide crRNA-alone effects against TuMV in *N. benthamiana*, but reported no impact on viral accumulation^16^. The report only included visible assessment, but no further molecular characterization such as quantifying the level of TuMV or confirming expression of the crRNA, and therefore the data are not conclusive, and the effect of GIGS may have gone unnoticed. Another report in *N. benthamiana* testing Cas13 variants also expressed guide-alone crRNA targeting the tobacco mosaic virus and no GIGS phenotype was reported ^18^. The experiment did not include data confirming expression of the crRNA, which could explain the difference, or the discrepancy may be due to other technical differences.

An important distinction for the experiments reported here, is our use of multi-guide crRNA in the absence of Cas13. To our knowledge, this control has never been reported in any eukaryotic system to-date. Our results suggest that multi-guides in the absence of Cas13 produce substantially more target RNA reduction compared to single-guides alone. Further research is needed to replicate this effect and understand why targeting discontinuous regions produce significantly more RNA reduction. Our extensive characterization of the GIGS phenomena in *N. benthamiana*, demonstration in tomato, verification in stably transformed *A. thaliana*, and evidence provided for a Cas9 designed crRNA, collectively show that guides cause target mRNA reduction on their own. Our results in plants are also consistent with the report of Cas13-independent transcript silencing in mosquito^43^. We posit that the findings described in mosquito represent the same GIGS phenomena reported here, which suggests that GIGS functions broadly across eukaryotes.

We found that GIGS elicits the production of sRNA with sequence corresponding to the targeted mRNA. Interestingly, multi-guide crRNA stimulated more sRNA production than single-guides, with the majority of sRNA corresponding to the crRNA target sequence, but we also found secondary sRNA targeting intervening regions with sequence not expressed in the crRNA. Given that sRNA are a hallmark of RNAi, it is likely that GIGS functions through endogenous components of RNAi. Further supporting this hypothesis, we found that sequence mismatches at positions 10,11 relative to the 5’ crRNA guide sequence, abolished the observed GIGS phenotypes and nearly eliminated target mRNA reduction. We infer these results to show that GIGS is dependent on the endonuclease activity of Argonaute. Interestingly, for Cas13 based crRNA to associate with AGO, it is likely they would first require processing. One possibility for the biogenesis of siRNA from a crRNA could be the processing of the crRNA-mRNA duplex. This could be carried out by one or more Dicer or Dicer-like endogenous ribonuclease III (RNase III) enzyme(s)^46^. While Dicer is conserved across eukaryotes, the gene family has differentially expanded, with a single copy present in vertebrates, two copies present in insects, and up to four Dicers in plants ^47,48^. It is possible that the duplication and diversification of the Dicer superfamily across eukaryotes will affect their competence for GIGS. Differences in Dicer substrate processing have been documented in eukaryotes^49,50^, and further mechanistic understanding is needed for multi-guide crRNA-mRNA processing. Aside from GIGS, it will also be important to determine if Cas13-mediated mRNA cleavage products interact with RNAi machinery to create feedback between the two RNA degradation systems.

The work presented here suggests that GIGS can achieve target RNA silencing using a guide sequence that is shorter than conventional hairpin and antisense constructs used in plants 51,52. This property could be particularly helpful in constructing compact multigene silencing cassettes expressed as a single transcript, which would significantly expand the capabilities of user defined RNA reduction schemes. In principle, multi-guide multi-target silencing could afford a higher target specificity compared to multi-gene RNAi given the significantly shorter expressed sequences, while avoiding the need to express a Cas13 transgene. Thus, GIGS based transcriptome engineering could provide a flexible *cis*-genic approach for plant biotechnology.

## Acknowledgments

We thank Bart PHJ Thomma for providing helpful comments during preparation of this manuscript. This work was supported by the Defense Advanced Research Projects Agency (grant no. D17AP00034) to D.E.C. The funder had no role in the study design, data collection and analysis, decision to publish, or preparation of the manuscript.

## Data Accessibility

Original and processed files for the small RNA sequencing data described in this research have been deposited in NCBI’s Gene Expression Omnibus (GEO)^53^, and are accessible through GEO Series accession number GSE171980, also accessible through BioProject PRJNA721612. (https://www.ncbi.nlm.nih.gov/geo/query/acc.cgi?acc=GSE171980).

## Author Contributions

D.E.C conceived the project. V.K.S., S.M., W.G.Z., D.M., G.P.M. and D.E.C. designed the experiments. V.K.S., S.M., W.G.Z., D.M., J.H., and W.Z. performed the experiments. V.K.S., S.M., W.G.Z., D.M., G.P.M. and D.E.C. analyzed the experiments. All authors contributed to writing the manuscript.

## Supplementary Data

Figure S1. Cas13a mediated efficient virus interference.

Figure S2. CRISPR inhibits TuMV with and without the Cas13 protein.

Figure S3. GIGS can function systemically to achieve virus interference.

Figure S4. Guide crRNA design and target sites for endogenous mRNA reduction by GIGS.

Figure S5. Endogenous mRNA reduction mediated by Cas13-dependent and GIGS expression.

Figure S6. Guide targets and experimental design for systemic endogenous mRNA reduction by GIGS.

Figure S7. Systemic endogenous mRNA reduction by GIGS.

Figure S8. Cas13-dependent and GIGS T1 transformed *A. thaliana* lines display phenotypes consistent with *TTG1* reduction.

Figure S9. Expression and translation products of Cas13 transgenic *Arabidopsis.*

Figure S10. Guide crRNA with mismatches at base pairs 10,11 do not elicit GIGS. Figure S11. Cas9 sgRNA can elicit GIGS photobleaching in *N. benthamiana*.

Table S1. Plasmids and gene sequences

Table S2. Backbones for cloning and expression of crRNA

Table S3. crRNA sequences for targeting of TuMV

Table S4. crRNA sequences for targeting of *Nicotiana benthamiana PDS*

Table S5. crRNA sequences for targeting of tomato *PDS*

Table S6. Oligo sequences used in this study

Table S7. Probe sequences for Nano-counting

Table S8. crRNA sequences for targeting of *Arabidopsis TTG1*

## MATERIALS AND METHODS

### Designing CRISPR-Cas13a machinery for *in planta* expression

To develop prokaryotic CRISPR-Cas13a machinery as a platform for *in planta* transcript-silencing, sequences of LbuCas13a and LbaCas13a effectors were *N. benthamiana* codon optimized along with 3x-FLAG tag or 3x-HA tag at the N-terminus, and custom synthesized (Genscript, Piscataway, NJ) (Supplementary Table S1). These fragments were assembled using HiFi DNA assembly (New England Biolabs, Ipswich, MA). The integrity of the constructs was confirmed by Sanger sequencing (Genewiz, South Plainfield, NJ).

Turnip mosaic virus engineered to express GFP (TuMV-GFP)^20^ and the endogenous phytoene desaturase (*PDS*) gene were selected as targets for CRISPR-Cas13a interference. For crRNA designs, Lba- or LbuCas13a specific direct repeats with 28 nucleotide spacer sequences complementary to the target were expressed by the *Arabidopsis thaliana* U6 promoter (Supplementary Table S2). For TuMV targeting, three single crRNAs targeting different regions of TuMV namely 5’untranslated region (5’ UTR), Helper component Proteinase (HcPro), viral genome linked protein (Vpg), and a poly crRNA containing aforementioned individual crRNAs in an array were designed and constructed (Fig. 1 and Supplementary Table S3). Similar to TuMV, the *PDS* transcript was targeted using three single crRNAs namely, s-guide 1, s-guide 2, and s-guide 3 and a multi-guide crRNA containing the three single guides (Supplementary Tables S3 and S4). To create mismatch guides corresponding to *PDS* multi-guide crRNA, the nucleotide sequence was altered at positions 5-6 bp, 10-11bp, and 21-22 bp from the 5’ end of each crRNA (Supplementary Table S4). A non-targeting crRNA was designed as a negative control. To create the sgRNA2 construct, we assembled the single-guide 2 target sequence with the transactivating crRNA (tracrRNA). The same strategy was used to construct sgRNA2 [50%mm] in which single-guide 2 crRNA had mismatches at every-other nucleotide. The NT-sgRNA negative control contained the Cas9 tracrRNA sequence and a non-plant target sequence (Supplementary Table S4).

### Cloning of CRISPR-Cas13a machinery

A backbone harboring AtU6 promoter sequence with one Lbu or Lba specific direct repeat sequence and *Bsa*I Golden Gate site was custom synthesized (IDT, Coralville, IA) for expressing crRNAs. This backbone was cloned into entry vector *pENTR* (Thermo Scientific, Waltham MA) using Topo cloning. Spacer sequences were ordered as oligos and cloned using *Bsa*I Golden Gate site. Gateway assembly (Invitrogen) was used to clone the promoter and crRNA cassette into the destination vector *pGWB413* containing or lacking Cas13a effector (Supplementary Table S1).

### Cloning crRNA for TRV systemic delivery

For systemic expression of crRNA using TRV, pea early browning virus (PEBV) promoter sequence with LbuCas13a specific direct repeat and *Bsa*I Golden gate site were custom synthesized (IDT, Coralville, IA) and cloned into Gateway entry vector *PCR8* (Supplementary Table S1). Three single guide and multi-guide crRNA sequences targeting *NbPDS*, and a multi-guide crRNA targeting *SlPDS* were ordered as oligos and cloned using Golden gate assembly (Supplementary Table S5). The cassette harboring PEBV promoter and TuMV, *NbPDS, or SlPDS* targeting crRNAs was PCR amplified with primers having *EcoR*I and *Mlu*I restriction sites and cloned into *EcoR*I and *Mlu*I digested *pTRV2* vector (Supplementary Table 6).

### Cloning of intron hairpin RNAi (hpRNAi) cassette

For cloning of PDS hpRNAi construct, a 197 bp sequence of PDS gene was custom synthesized as sense and antisense arm along with PDK intron sequence with 25 bp overhang complementarity to *pGWB413* vector (Supplementary Table S1). All the fragments were assembled using HiFi DNA assembly (New England Biolabs, Ipswich, MA) expressed by the 35S promoter.

### Agro-infiltration of *N. benthamiana* and *Solanum lycopersicum*

*N. benthamiana* plants were grown and maintained in growth chamber at 23°C with 16-hour day and 8 hour light cycle and 70% humidity. Four-week-old plants were used for leaf spot agroinfiltration to test Cas13a interference against TuMV-GFP. Binary constructs harboring Cas13a homologs with or without crRNA (targeting TuMV or *PDS* transcript), TuMV-GFP infectious clone (a gift from Dr. James Carrington) were individually transformed into chemically competent *Agrobacterium tumefaciens* strain GV3101. Single colonies for each construct were inoculated into LB medium with antibiotics and grown overnight at 28 °C. Next day, the cultures were centrifuged and suspended in agroinfiltration buffer (10mM MgCl2, 10mM MES buffer pH 5.7 and 100μM acetosyringone), and incubated at ambient temperature for 2-3 hours. For TuMV interference assay, *Agrobacterium* cells harboring Cas13a with crRNA targeting TuMV were infiltrated at an OD600 of 1.0 into adaxial side of four-week-old *N. benthamiana* leaves using a 1.0 ml needleless syringe. Two days later, *Agrobacterium* cells harboring TuMV-GFP were infiltrated into same areas at an OD600 of 0.3. After five days, interference activity of Cas13a against the TuMV-GFP was assayed by visualizing GFP in infiltrated leaves under UV light using a hand-held UV lamp (Fisher Scientific, Waltham, MA) and a Nikon camera.

For PDS silencing, leaves of four-week-old *N. benthamiana* plants were infiltrated with *Agrobacterium* cultures harboring LbuCas13a with crRNAs targeting PDS and leaf samples were collected at 5 days post inoculation. For TRV mediated crRNA delivery, assays used three-week-old *N. benthamiana* plants. A single colony of *Agrobacterium* harboring crRNAs targeting *PDS* were inoculated into LB medium with antibiotics and grown overnight at 28 °C. Next day, the cultures were centrifuged and resuspended into infiltration buffer at an OD600 of 0.6. The cultures were incubated at ambient temperature for 2-3 hours and infiltrated into *N. benthamiana*. Two upper leaves were collected two-weeks after TRV infiltration. Control plant infiltrated with TRV expressing an RNAi antisense fragment were used to help track systemic TRV movement. Infiltration of tomato plants was performed similarly to *N. benthamiana* except that *Agrobacterium* cells were resuspended into infiltration buffer at an OD600 0f 2.0. The cultures were incubated at ambient temperature for 2-3 hours and infiltrated into three-week-old tomato plants. Data was collected two-weeks after TRV infiltration in the lower leaves.

### RNA isolation, cDNA synthesis, qRT-PCR and northern blotting

Total RNA was isolated from Agro-infiltrated leaf samples and upper leaf tissue following systemic TRV movement using Trizol (Ambion) ^54^. For first strand cDNA synthesis, DNase treated 1 μg total RNA was reverse transcribed using either random hexamers or oligo(dT20) and SuperScript II reverse transcriptase (Thermo Fisher Scientific) according to the manufacturer’s instructions. Quantitative PCR was performed using SYBR Select Master Mix (Applied Biosystem) and gene specific primers (Supplementary table) for *PDS* and TuMV. *EF1*α gene was used as internal house-keeping reference for PDS and TuMV qRT-PCR ^55^ The experiments were repeated three times with three biological and two technical replicates. Relative expression values were plotted using ggplot2 in R ^56,57^. For detection of *PDS* transcript, 20 μg of total RNA was separated on a denaturing 1.2% agarose gel and blotted on a Hybond-N+ (Roche) membrane. RNA was crosslinked using UV light and hybridized with a DIG labelled probe (PCR DIG probe synthesis kit, Sigma). For detection of LbuCas13a the membrane was stripped and probed with DIG labelled Cas13a specific probe and signal detected on a Licor Odyssey imaging system (LI-COR Bioscience, Lincoln, NE).

### Real time quantification of PDS and TuMV transcripts using Nanocounting technology

For direct RNA quantification of PDS and TuMV transcripts using NanoString technology, we collected sequence data for different *N. benthamiana* genes including *PDS*, three house-keeping genes for normalization (*PP2aa2*, *EF1α*, *RPL23a*), LbuCas13a, HCPro and coat protein (Supplementary Table 7). The sequence information was utilized to design two probes for each target gene. Total RNA samples (300 ng total RNA) and probe master mix were supplied to the Huntsman Cancer Institute, University of Utah for Nanostring quantification following manufacturer specifications. The nano-counting data was analyzed using the nSolver software.

### Western blotting

For western blotting, total protein was isolated from *Agrobacterium* infiltrated leaves using extraction buffer (50mM Tris-Cl, 1% β-Mercaptoethanol and protease inhibitor cocktail (Roche, Basel, Switzerland)). Total proteins were boiled with loading buffer (100mM Tris-Cl, 20% Glycerol, 4% SDS, 10% β-Mercaptoethanol and 0.2mg/ml bromophenol blue) and resolved on 12% SDS-PAGE gel. The proteins were transferred from SDS-PAGE gel to PVDF membrane (GE healthcare, Chicago, IL). Membrane blocking and antibody incubations were performed using iBind western device (Thermo Fisher Scientific, Waltham, MA) according to the instrument manual. Finally, the membrane was treated with ECL Select western blotting detection reagent (GE healthcare, Chicago, IL) and signal was detected with Licor Odyssey imaging system (LI-COR Bioscience, Lincoln, NE).

### Small RNA sequencing and analysis

Two separate small RNA sequencing experiments were conducted. For results shown in (Fig. 3a-e), Cas13 and crRNA guides and controls were expressed in *N. benthamiana* leaves using agrobacterium spot infiltration as described. Total RNA was extracted from infiltrated leaves using Trizol following manufactures guidelines. For results shown in (Fig. 3f-l), crRNA guides and controls were expressed from TRV using agrobacterium infiltration as described. Total RNA was extracted from upper leaves following systemic TRV movement using Trizol. Total RNA samples were sent to the Beijing Genomics Institute (BGI Group, Hong Kong). Twenty-four small RNA libraries were constructed following the DNBseq small RNA library protocol. Briefly, small RNA were isolated from PAGE gel corresponding to size 18-30 nt. Adapters were ligated and first strand synthesis performed according to DNBseq small RNA library protocol. Libraries were PCR amplified and size selected and sequenced on the DNBseq platform (BGI Tech Solutions, Hong Kong, China).

Small RNA reads for both experiments were trimmed ^58,59^, and aligned using STAR (v2.7.3a) ^60^ to a modified version of the *N. benthamiana* genome (v1.0.1)^61^. The modifications included removing all contigs with less than 70K nt, adding the coding sequence of LbuCas13a as a contig, and masking one of the two paralogs coding for PDS. The coding sequence for *PDS* on contig Niben101Scf14708, position 12885-21779 (gene23) was masked in order to ensure unique mapping to a single *PDS* locus on contig Niben101Scf01283, position 197129-205076 (gene 2002). Uniquely mapped read counts for the exons were extracted per base-pair using samtools (v1.3)^62^ and bedtools ‘coverage’ (v2.29.2) ^63^. To compare between sequenced samples, mapped reads were normalized to library size (i.e. total uniquely mapped reads per library) using the equation (number of reads mapped at a nucleotide position * (1 / number of uniquely mapped reads in library) * 1M), referred to as counts per million (CPM). The size distribution of uniquely mapped reads were analyzed for 21, 22, and 24 nt sRNA. The average number of uniquely mapped sRNA to the PDS transcript was calculated for the duplicate samples for each size class. The proportion of each size class was determined by the equation, ((average number of reads per size class / sum of average number of reads per size class)*100). Analyses were carried out using Python3 (v3.8.2) libraries NumPy (v1.18.1), Pandas (1.0.3) and plotted with Matplotlib (v3.2.1) ^64–67^. Processed files, additional information and the reference genome used for mapping are provided through the GEO^53^ Series accession number GSE171980. (https://www.ncbi.nlm.nih.gov/geo/query/acc.cgi?acc=GSE171980).

### Generating stable transgenic Arabidopsis plants

*TTG1*-targeting three single guides (guide-1, −2, −3) and a multi-guide crRNA (Supplementary Table 8), and non-targeting (NT) oligos were annealed and ligated into *pENTR* backbone containing *Bsa*I Golden gate site. Gateway assembly was used to transfer guide crRNA to *pGWB413* destination vector with or without 3xHA-LbuCas13a. Stable transgenic *Arabidopsis* plants expressing *TTG1* guides with or without LbuCas13a were generated using *Agrobacterium*-mediated floral dip ^68^ Similarly, stable *Arabidopsis* controls with a NT crRNA, a 197 bp hairpin construct against *TTG1* (a gift from Dr. Steven Strauss), and no guide transformation control (only 3xFLAG-LbuCas13a) were generated. One month after floral dip, T_1_ seeds were collected and stored at 4°C.

### Arabidopsis phenotyping

Transformed T_1_ Arabidopsis seedlings were identified using rapid selection protocol ^69^. Selection was conducted on ½ MS media with a Kanamycin concentration of 100 μg/ml. Positive transformants (*n* = 36) for each *TTG1* crRNA with or without LbuCas13a and *TTG1* hairpin controls were transferred to soil and grown under optimal conditions. Control Arabidopsis Col-0 plants were germinated on ½ MS media without Kanamycin and transferred to soil. Seventh leaf from ten individual plants for each construct was imaged under a dissecting microscope equipped with a Nikon camera and trichomes were counted using multi-point feature in ImageJ software ^70^. For each construct, RNA was extracted from 10^th^ leaf of five individual plants with varying leaf trichomes to quantify *TTG1* expression using qRT-PCR. *AtEF1α* was used as internal house-keeping control for normalizing *TTG1* expression (Supplementary Table 6). Selected individual plants for each construct were self-pollinated to collect T_2_ seed. Five technical replicates of each selected plant/line were used for analyzing total flavonoids, in 5 mg seed, using modified aluminum chloride (AlCl_3_) colorimetric method ^71^. Total flavonoids content was estimated using the following formula: flavonoids (mg/g) = concentration obtained through quercetin calibration curve × (volume of extract/seed weight).

To determine the inheritance of GIGS and Cas13-mediated gene silencing, 10 T_2_ plants from selected T_1_ lines were transferred to soil after Kanamycin selection. Seventh leaf from 10 individual T_2_ plants was imaged for counting leaf trichomes. Statistical comparisons between the transformation control (no guide) and each selected line was performed. *TTG1* expression in the top rosette leaf from three individual T_2_ plants was analyzed using qRT-PCR. Five individual T_2_ plants for each line were self-pollinated to collect T_3_ seed. Total flavonoid content was analyzed in T_3_ seeds from five independent seed lots (five biological replicates). Similarly, proanthocyanidins content was measured using DMACA-HCl method from three seed lots ^72^. Proanthocyanidins were measured at 640 nm and reported as per gram of seed weight. Total flavonoid and proanthocyanidin analyses were repeated twice, the averaged values for each seed lot were used for statistical comparisons. Absorbance of flavonoids and anthocyanin was measured using Thermo Spectronic 3 UV-Visible Spectrophotometer. While absorbance of proanthocyanidins was measured through Synergy H1 Hybrid Multi-Mode Microplate Reader (Agilent Technologies, Winooski, Vermont).

For leaf anthocyanin quantification, one-week-old T_3_ seedlings after Kanamycin selection were transferred into ½ MS media + 3% sucrose and subjected to light stress (500 μmol m^−2^ s^−1^) for one week. 200 mg of leaf tissue was used for quantifying anthocyanin ^73^. Anthocyanin analysis was repeated twice with 5 replicates in each batch. Anthocyanin content was calculated by using following formula (absorbance/35,000× dilution factor×647 × 1,000 per mg of sample extracted (in mg g-1 fresh weight). Representative plantlets following sucrose treatment showing anthocyanin pigmentation were imaged with a dissecting microscope equipped with a Nikon camera. To test *TTG1* expression in T_3_ generation, seventh leaf from three individual plants was analyzed using qRT-PCR. To determine the expression of LbuCas13a, RT-PCR was conducted on cDNA synthesized for qRT-PCR. Western blot analysis with HA-tag antibody was conducted on one-week-old T_3_ seedlings post Kanamycin selection.

