## Supplemental Figures for "RNA silencing by CRISPR in plants does not require Cas13"

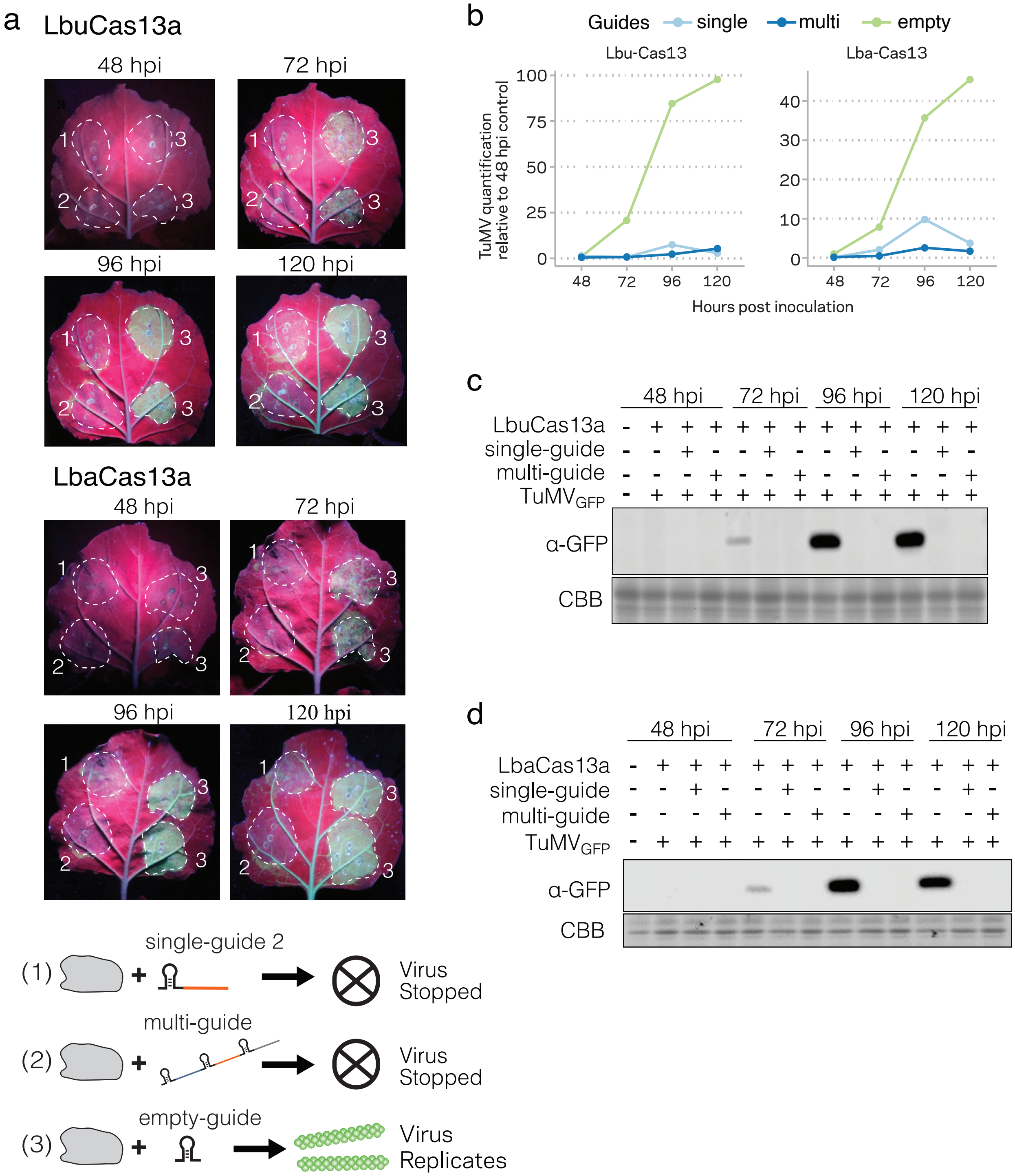


**Supplementary Figure 1. Cas13a mediates efficient virus interference. a,** Images of *N. benthamiana* leaves from 48 hours post inoculation (hpi) to 120 hpi shown under UV light to visualize GFP fluorescence. The Cas13 protein (either LbuCas13 or LbaCas13) were expressed with a guide and the TuMV expressing GFP virus as indicated by the schematic diagram and numbering. The areas of agro-infiltration are indicated with white dashed circles. Higher GFP signal results from increased virus accumulation **b,** Quantification of viral genome accumulation using qPCR. TuMV levels were standardized to plant endogenous *EF1α* transcript and three samples were collected per time point. **c,d,** GFP protein accumulation over the time series of inoculated leaves expressing LbuCas13a (c) or LbaCas13a (d). The lanes are labeled above the images for each time point, while the anti-GFP (α–GFP) panel shows signal from western blot, and the commassie brilliant blue (CBB) stained gel panel shows the loading control.

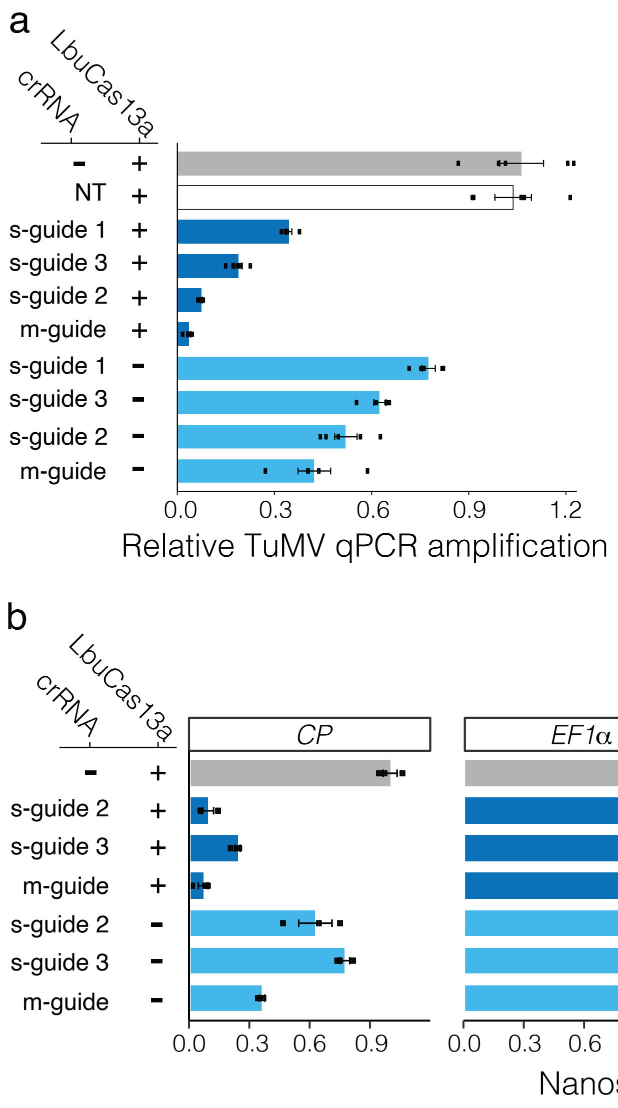


**Supplementary Figure 2. CRISPR-Cas13a inhibits TuMV with and without the Cas13 protein. a,** Quantification of TuMV accumulation from *N. benthamina* transient spot expression. Leaves were inoculated with TuMV and a combination of Cas13 and guide crRNA as indicated to the left of the barplot. Individual samples are shown as black dots and the mean is shown as a bar with standard error. TuMV levels were standardized to the plant endogenous *EF1α* transcript and normalized to the Cas13 alone sample. Five samples were collected for each treatment. **b,** Nanostring quantification for using four different probes: TuMV (Coat protein, CP), and three endogenous controls from *N. benthamiana* (*EF1a*, *PP2aa2*, *RPL23a*). Samples expressed either the LbuCas13a protein (+) or no transgene (-), along with no-guide (-), a single-guide (s-guide), or a multi-guide crRNA (m-guide). Three independent samples were analyzed per treatment.

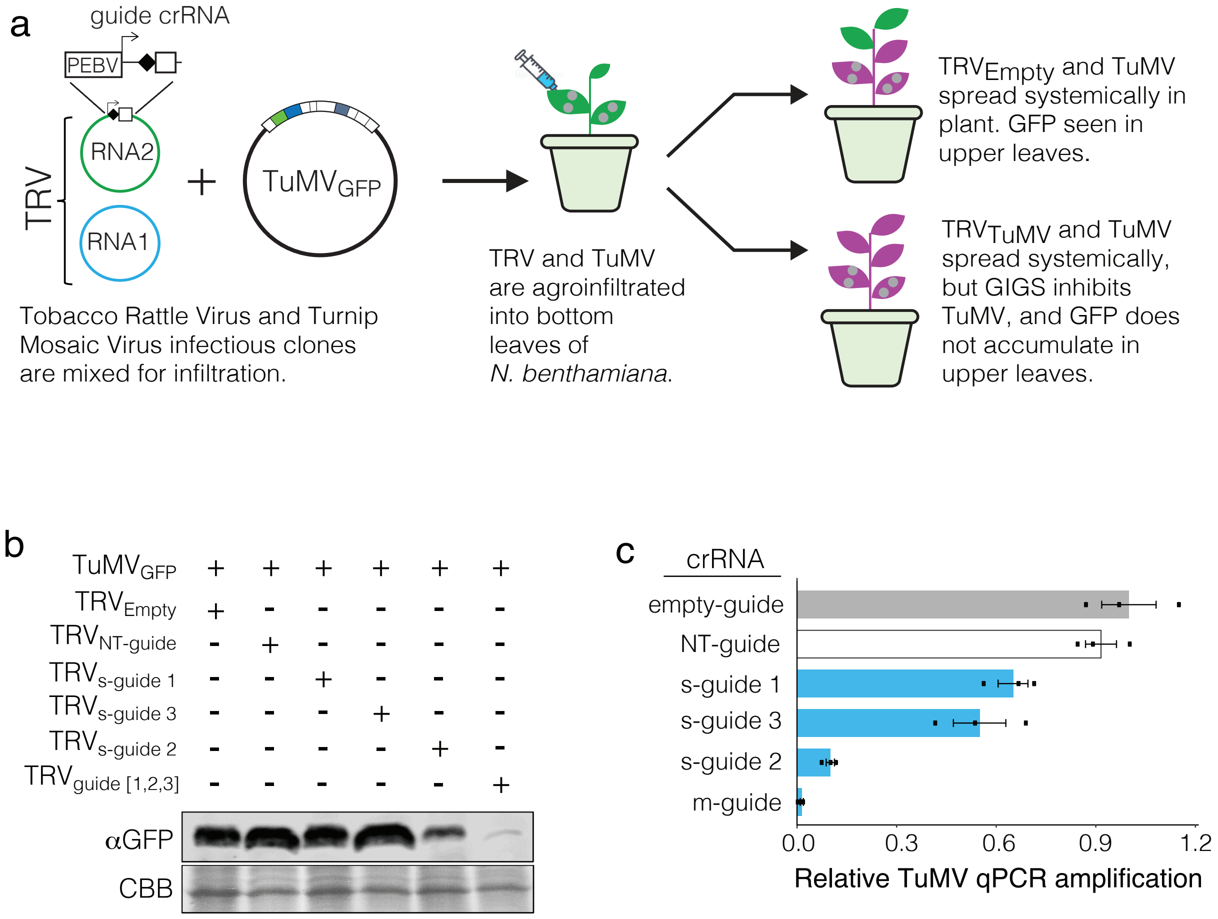


**Supplementary Figure 3. GIGS can function systemically to acheive virus interference. a,** Schematic diagram of the TRV expressing guide crRNAs and TuMV expressing GFP. The two infectious clones are mixed and agroinfiltrated into *N. benthamiana*. At 7 days post inoculation, plants are visualized under ultraviolet light. TRV expressing an empty or NT-guide does not target TuMV and GFP accumulates in upper leaves. TRV expressing crRNA targeting TuMV results in GIGS and the plants fail to accumulate GFP. **b,** Upper leaves were checked for GFP accumulation by western blot corresponding to images shown in (Fig. 1d). The GFP antibody panel (αGFP) shows GFP signal while commassie brilliant blue panel (CBB) shows protein loading. **c,** The TuMV genome was quantified from three independent plants (shown as black points) using qPCR. The mean and standard deviation are shown as barplots. Samples were standardized to the *EF1α* endogenous transcript and normalized to the empty-guide control levels.

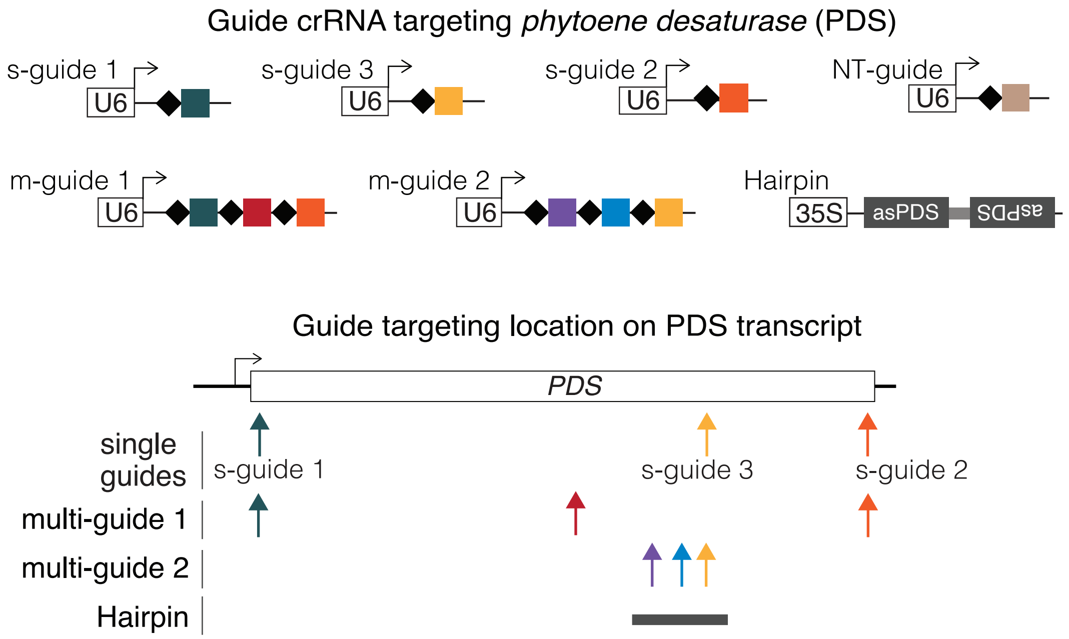


**Supplementary Figure 4. Guide crRNA design and target sites for endogenous mRNA reduction by GIGS.**

**a,** Schematic of single- and multi-guide crRNA (s-guide and m-guide, respectively), along with control non-targeting guides (NT-guide) and RNAi inducing Hairpin. The approximate location where the respective guides are antisense to the PDS transcript (i.e. thier target location) are denoted by filled arrows. The region covered by the hairpin construct is shown as a grey bar.

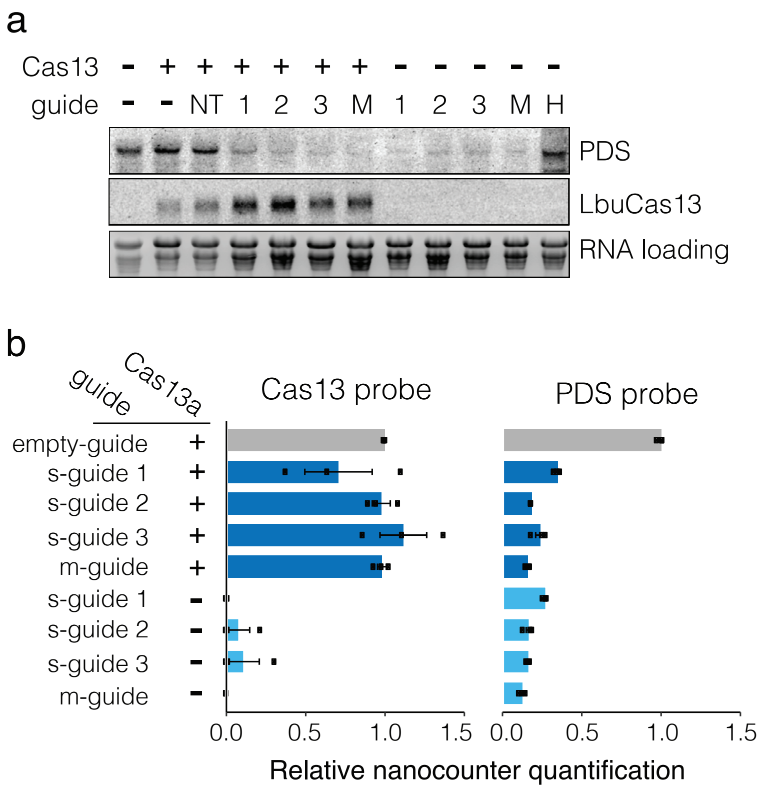


**Supplementary Figure 5. Endogenous mRNA reduction mediated by Cas13-dependent and GIGS expression.**

**a,** Reduction in *PDS* mRNA accumulation as measured by northern blot analysis using a PDS probe (PDS panel). Samples expressing the Cas13a transcript are indicated (Cas13 +) and correspond to signal from the Cas13 probe (LbuCas13 panel). Samples expressed no guide (-), a non-target guide (NT), one of three single-guides (1,2, or 3), a multi-guide (M), or a hairpin construct (H). The presence of roughly equal RNA amounts was confirmed by the abundance of ribosomal RNA signal (RNA loading panel). **b,** Quantification directly on RNA samples using nanostring for Cas13 (Cas13 probe) and *phytoene desaturase* (PDS probe) mRNA. The presence (+) or absence (-) of Cas13 expression is indicated to the left, along with the expression of a single-guide (s-guide) or the multi-guide (m-guide).

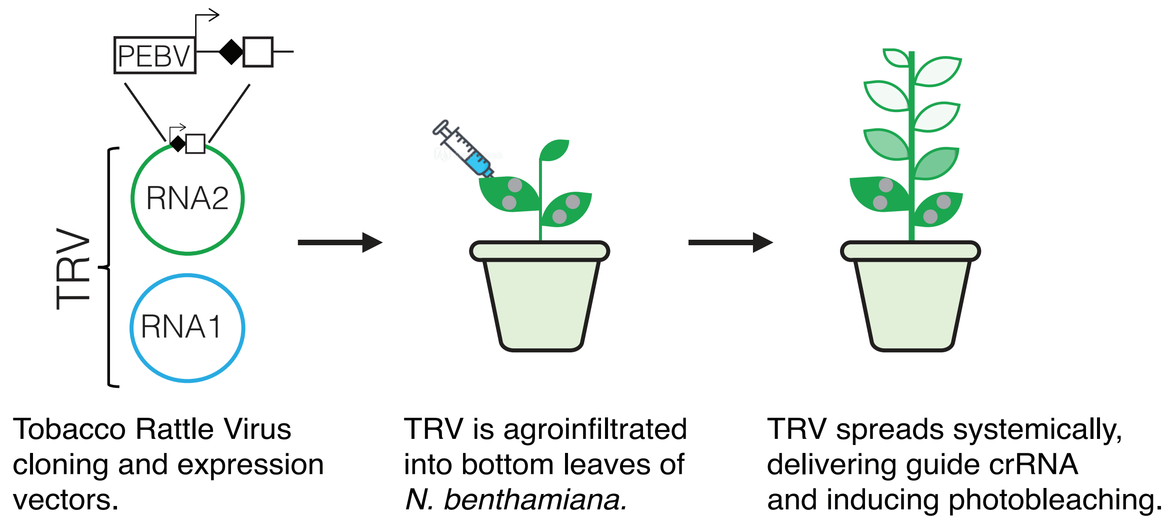


**Supplementary Figure 6. Guide targets and experimental design for systemic endogenous mRNA reduction by GIGS.**
Schematic of the TRV expression vector system (RNA1 and 2). RNA2 was engineered to contain the pea early browning virus (PEBV) promoter to express single-, multi-, and NT-guides. An antisense fragment (371 bp) to *PDS* was also inserted into the RNA2 cloning sight to induce RNAi against *PDS*. Infectious clones are agroinfiltrated. TRV moves systemically in the plant, delivering the respective guide crRNA. Those that cause *PDS* mRNA silencing result in a bleaching phenotype (i.e. white sectors) seen on upper leaves.

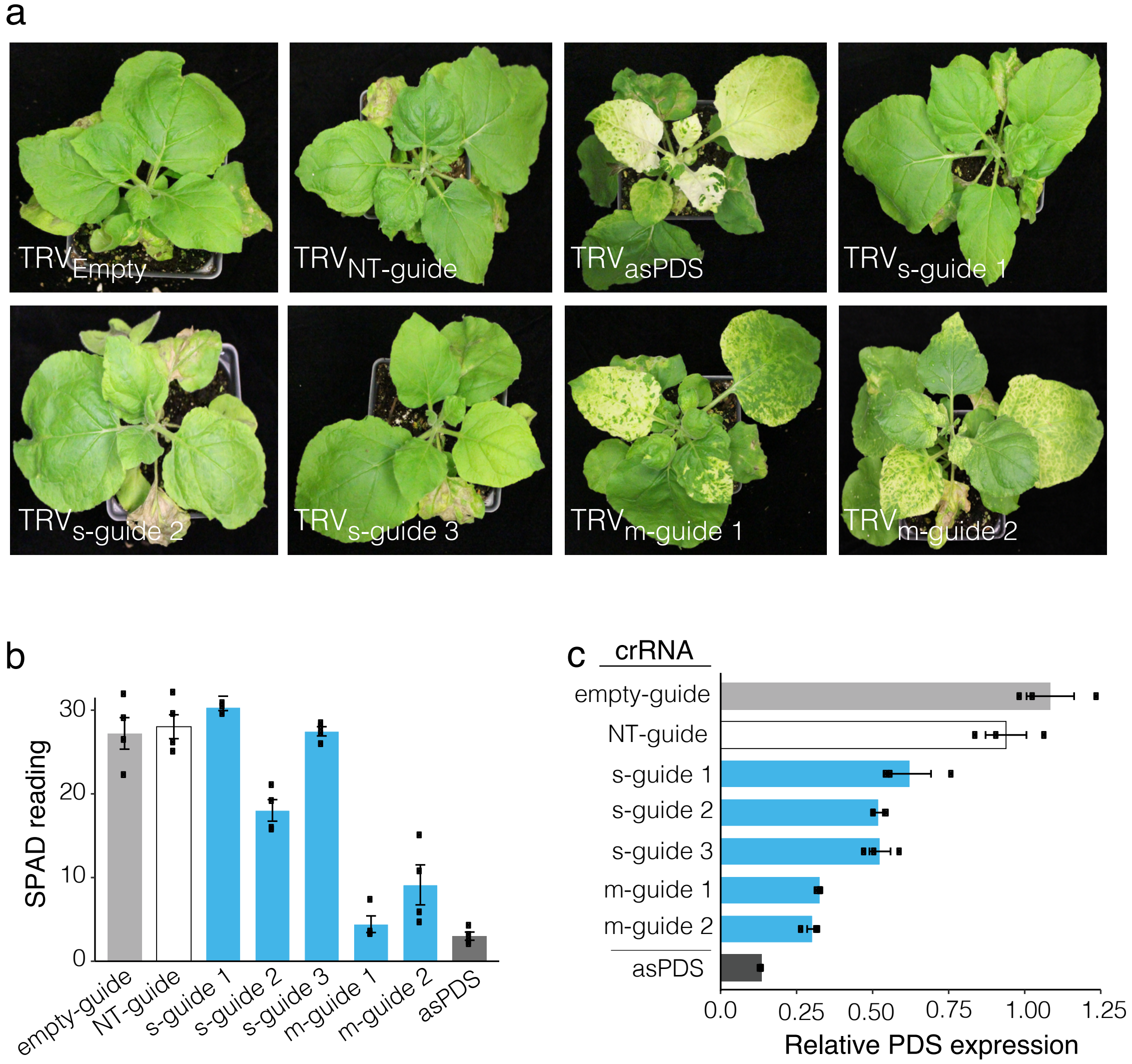


**Supplementary Figure 7. Systemic endogenous mRNA reduction by GIGS.**
**a,** Representative images of *N. benthamiana* plants two weeks post TRV inoculation. TRV moved systemically to the top portion of the plant and expressed guide crRNAs targeting *PDS* or controls. Photobleaching caused by *PDS* mRNA reduction is visible as white or yellow sectors in the upper leaves. Each image is labeled with the guide delivered by TRV. **b,** SPAD meter readings from photobleached areas of leaves. One reading was taken per infiltrated plant. Three independent plants were infiltrated. **c,** The *PDS* transcript was quantified using qPCR from three independent leaves. Data from each sample is shown as a black point with the mean and standard deviation shown as a boxplot. Samples were standardized to the *EF1*$\alpha$ endogenous transcript and normalized to the empty-guide control levels.

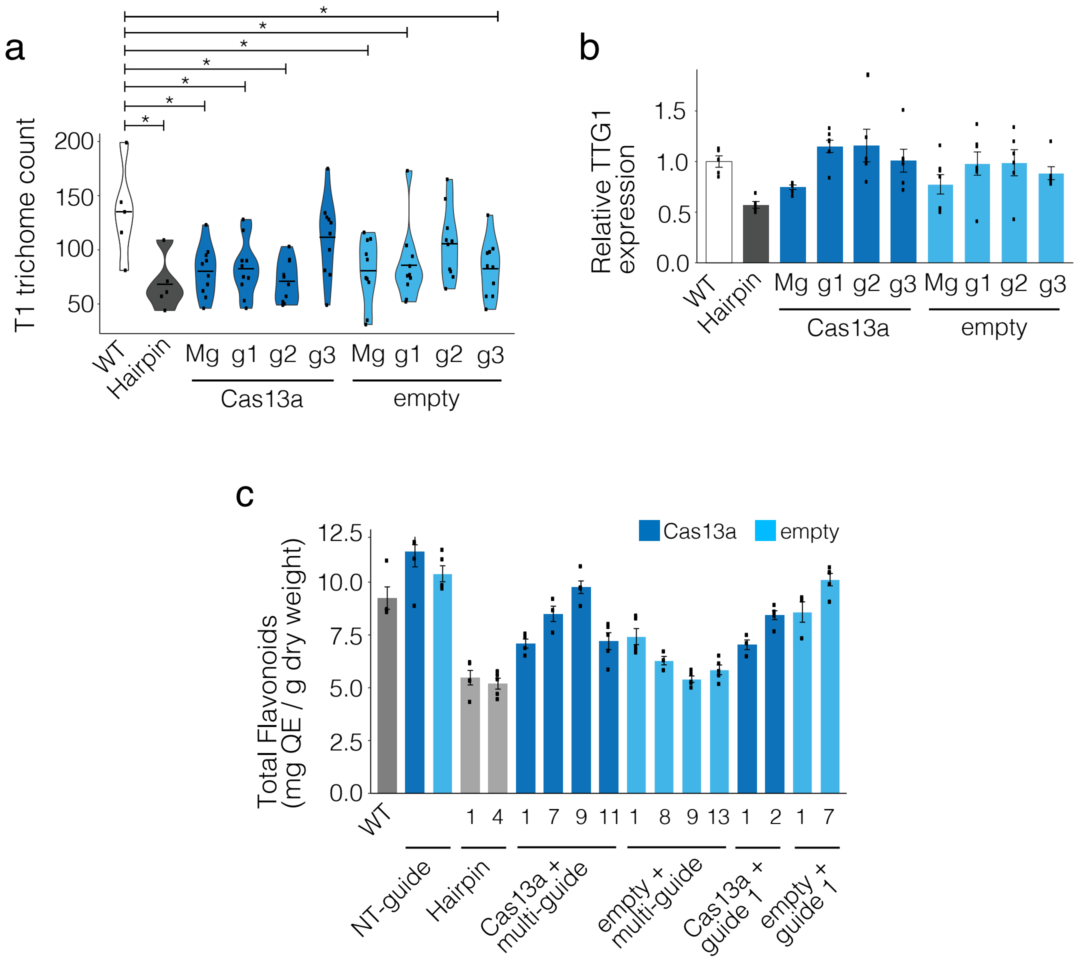


**Supplementary Figure 8. Cas13-dependent and GIGS $T_1$ transformed *A. thaliana* lines display phenotypes consistent with *TTG1* reduction.**

**a,** Leaf trichome counts from individual $T_1$ lines shown as black points and the distribution shown as a violin plot. Three single-guides (g1, g2, g3) or a multi-guide (Mg) were transformed into plants with Cas13a (dark blue) or without (light blue). Plants were transformed with a hairpin construct (grey) designed to silence the *TTG1* transcript. Statistical comparisons to the non-transformed control (WT, wild-type) were made as one-sided Mann-Whitney U-test with Benjamini-Hochberg (BH) multiple testing correction. Samples with p-values less than 0.05 (*) are indicated. **b,** Five lines from each transformation group were assessed for *TTG1* transript levels using qPCR. The values were standardized to *AtEF1*α endogenous control and normalized to the WT control. Individual data points are shown as black points and the mean and standard deviation shown as a barplot. **c,** Individual $T_1$ plants were self fertilized, and five lots of seed from each plant (technical replicates) were analyzed for seed total flavonoid content. Transformants expressing Cas13a are shown in dark blue while those not expressing Cas13a are shown in light blue. Individual data points are shown as black points and the mean and standard deviation shown as barplots. WT, wild-type; C13, Cas13a expressing; empty, no Cas13a protein; Hairpin, expressing a 197 bp hairpin against *TTG1*. Numbers indicate the line numbers for the indicated treatments shown below.

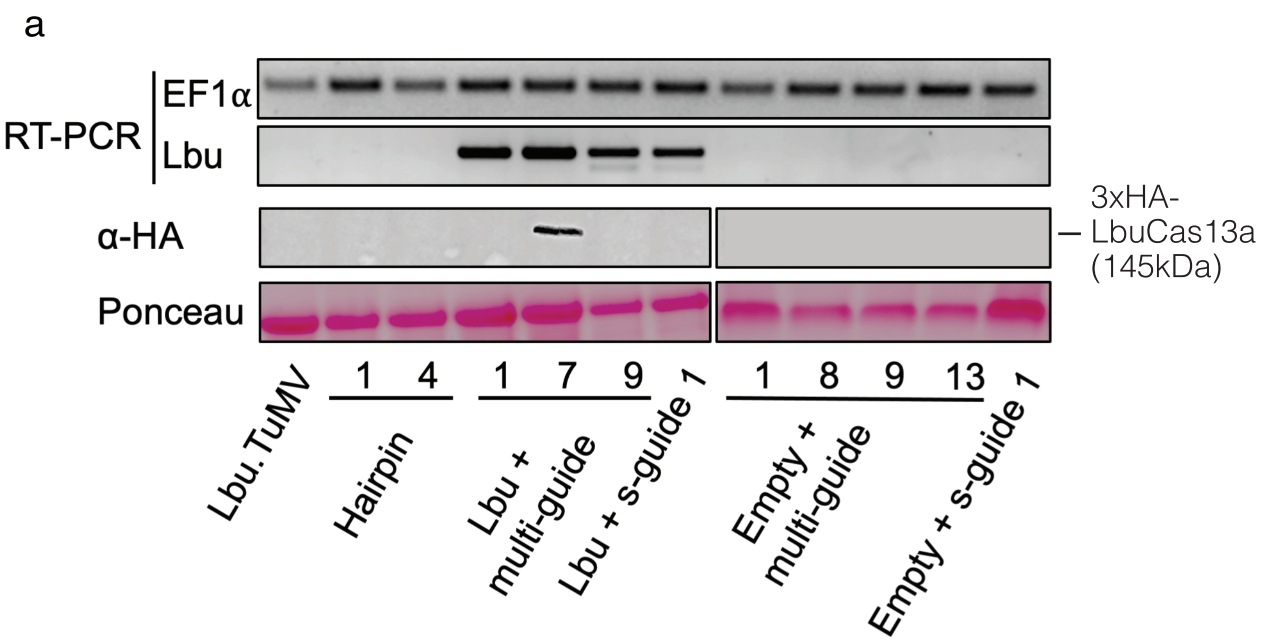


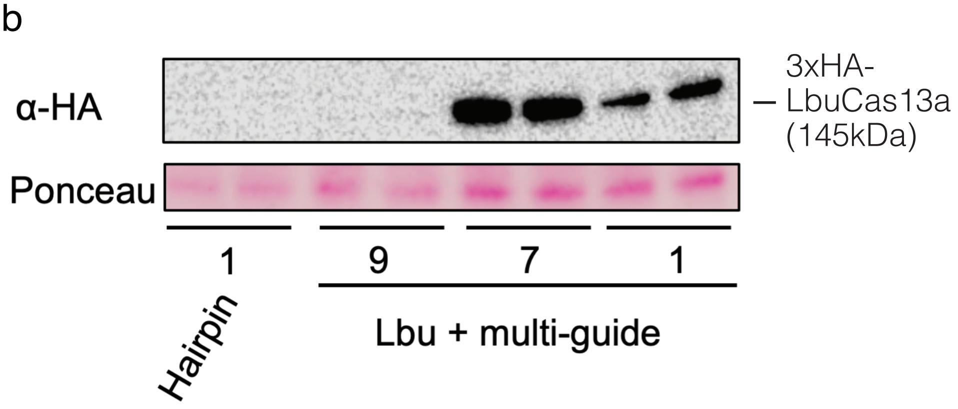


**Supplementary Figure 9. Expression and translation products for Cas13 targeting *TTG1* transgenic *Arabidopsis.***

**a**, The Cas13 transcript is only detected using reverse-transcription PCR (RT-PCR) in lines transformed with the LbuCas13a transgene (Lbu, top panel). The endogenous transcript coding for *EF1*α is shown as a control. The western blot panel using an anti-HA antibody (α-HA) detected a single band corresponding to the size of the 3xHA-LbuCas13a protein (145kDa). Ponceau strain was used a protein loading control.
**b**, Protein was isolated from the same lines used in (a) and re-analyzed by western blot. Higher exposure time detected a single band corresponding to 3xHA-LbuCas13a in line #1 as well as line #7. The results indicate likley differences in protein translation or possibly stability between the different transgenic lines.

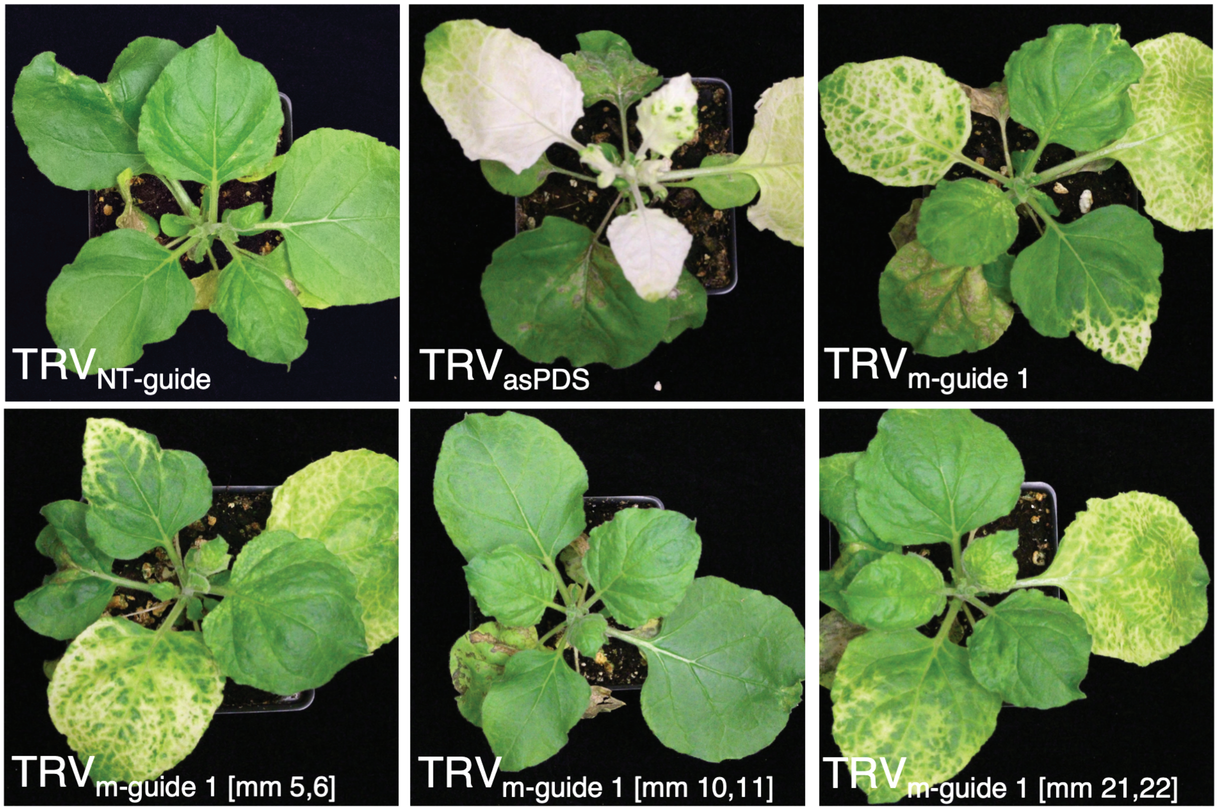


**Supplementary Figure 10. Guide crRNA with mismatches at base pairs 10,11 do not elicit GIGS.**

Representative images of plants following tobacco rattle virus (TRV) systemic movement. Plants infected with TRV expressing a non-targeting guide crRNA (NT-guide) have a normal green leaf appearance. Plants infected with TRV containing an antisense *PDS* fragment (asPDS) display photobleaching in upper leaves following TRV systemic movement and the triggering of RNAi *PDS* silencing. TRV expressing multi-guide 1 (m-guide 1), multi-guide 1 with mismatches between the target and guide at positions 5 and 6 (m-guide 1[mm 5,6]), or multi-guide 1 with mismatches at positions 21 and 22 (m-guide 1[mm 21,22]) also display photobleaching in upper leaves. Plants inoculated with TRV expressing multi-guide 1 with mismatches between the target and guide at positions 10 and 11 (m-guide 1[mm 10,11]) have the same normal green appearance as the NT-guide samples.

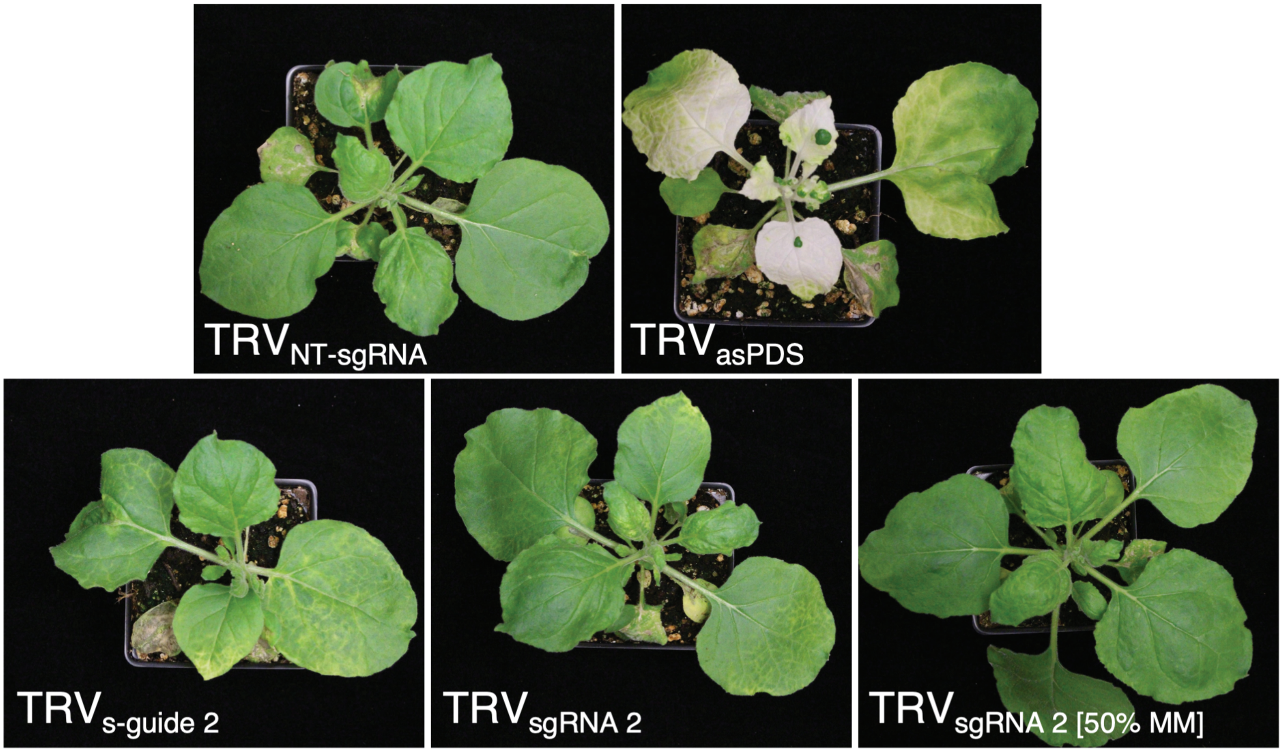


**Supplementary Figure 11. An sgRNA guide designed from the Cas9 system can elicit GIGS photobleaching in *N. benthamiana.***

Representative whole plant images showing GIGS induced photobleaching. Plants expressing the control non-targeting single guide RNA (NT-sgRNA) from TRV do not display abnormal leaf color. Strong photobleaching is seen from plants expressing TRV containing an antisense *PDS* fragment (asPDS). Clear phenotypic differences in leaf greenness are seen when TRV expressed s-guide 2 (crRNA designed based on Cas13 system) or sgRNA 2 (crRNA designed based on Cas9) targeting *PDS*. These plants show interveinal yellowing. The control Cas9 sgRNA 2 with 50% mismatches to the *PDS* transcript (sgRNA 2[50%mm]) does not display any visible alteration in leaf greenness.
